# Commensal-derived Trehalose Monocorynomycolate Triggers γδ T Cell-driven Protective Ocular Barrier Immunity

**DOI:** 10.1101/2025.03.17.643820

**Authors:** Xiaoyan Xu, Yannis E. Rigas, Mary J. Mattapallil, Jing Guo, Vijayaraj Nagarajan, Eric Bohrnsen, Crystal Richards, Akriti Gupta, Guillaume Gaud, Paul E. Love, Timothy Jiang, Amy Zhang, Biying Xu, Zixuan Peng, Yingyos Jittayasothorn, Mary Carr, M. Teresa Magone, Nathan T. Brandes, Jackie Shane, Benjamin Schwarz, Anthony J. St. Leger, Rachel R. Caspi

## Abstract

Commensals shape host physiology through molecular crosstalk with host receptors. Identifying specific microbial factors that causally influence host immunity is key to understanding homeostasis at the host-microbe interface and advancing microbial-based therapeutics. Here, we identify trehalose monocorynomycolate (TMCM) from *Corynebacterium mastitidis* (*C. mast*) as a potent stimulator of IL-17 production by γδ T cells at the ocular surface. Mechanistically, TMCM-driven IL-17 responses require both IL-1 signals and γδ TCR signaling, which also supports endogenous γδ T cell IL-1R1 expression. Notably, synthetic TMCM alone is sufficient to mimic the effect of *C. mast* in inducing γδ T cell immunity and protect against pathogenic corneal infection. Our findings establish TMCM as a key mediator of commensal-driven immune defense, highlighting its potential as a γδ T cell adjuvant and a microbiome-informed therapeutic to enhance IL-17-driven protection at barrier sites such as the ocular surface.

**Graphical Abstract:** 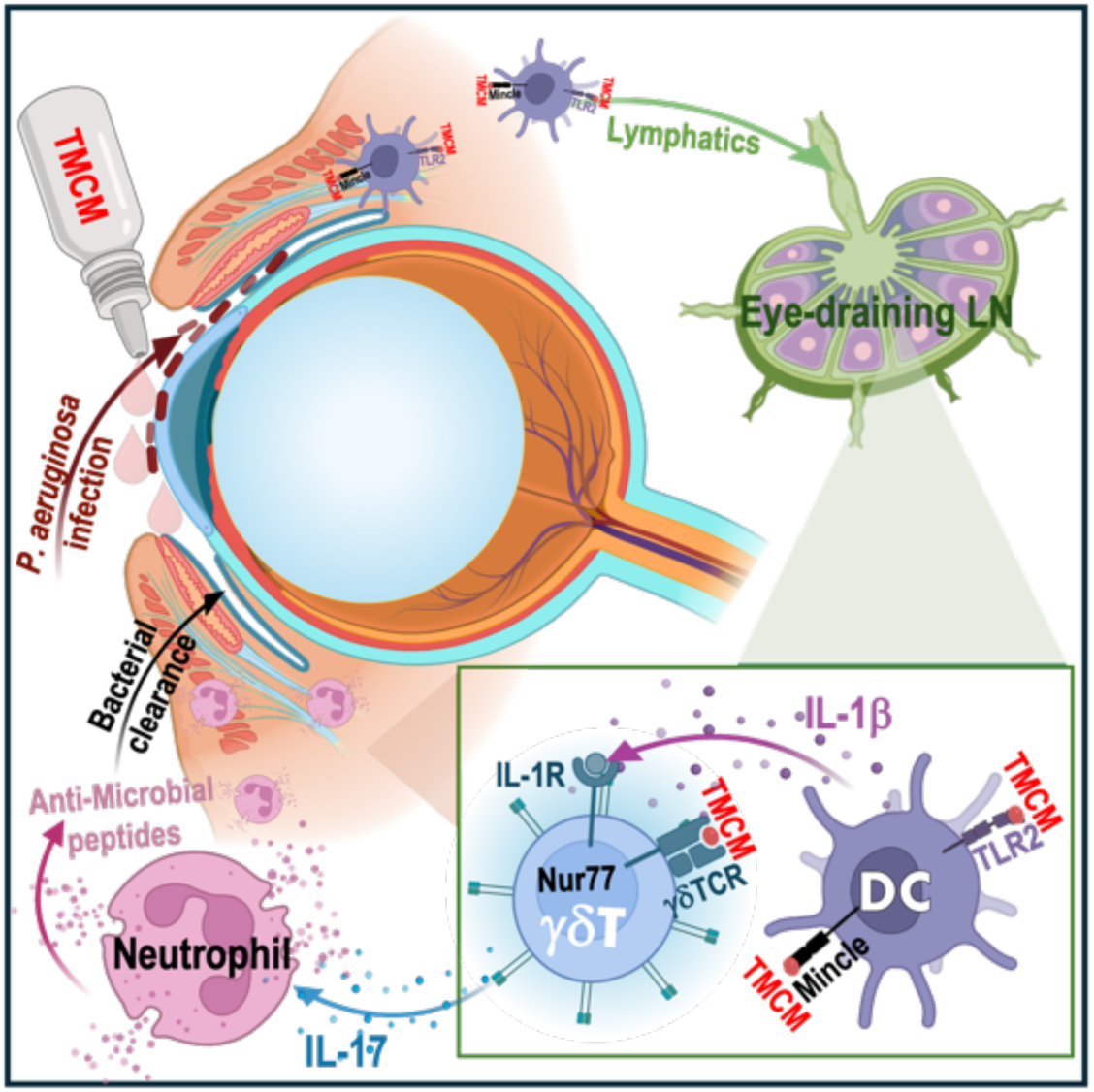

**HIGHLIGHTS:**

- Corynomycolates enable ocular *C. mast* colonization and protective IL-17 immunity
- TMCM drives IL-17 from γδ T cells through TCR and IL-1R signaling
- γδ TCR signaling maintains the expression of endogenous IL-1R1
- Synthetic TMCM mimics the ability of *C. mast* to induce γδ T cell immunity in the eye
- TMCM protects against *P. aeruginosa* keratitis, highlighting its therapeutic potential

## INTRODUCTION

The establishment of immune homeostasis between the host and commensal bacteria is essential for optimal tuning of host immunity to prevent infection^1,2^. Previous studies have shown that the host has evolved a variety of ways to recognize and respond to commensal bacteria, and the bacteria have evolved ways to survive attacks by the host immune system. These interactions are mediated mainly by molecular cross-communication between microbe-associated molecular patterns (MAMPs) from commensal bacteria and host pattern recognition receptors (PRRs) on innate immune cells, but TCR-driven interactions may also play a role^3^. Here, we identify a microbial factor, trehalose monocorynomycolate (TMCM), produced by, *Corynebacterium mastitidis* (*C. mast*), that stimulates γδ T cells, in part through the γδ TCR, and confers protection against ocular challenge with the pathogen, *Pseudomonas aeruginosa*.

Corynebacteria are commonly identified as core members of the microbiomes at the skin, in the oral/nasal cavity, and at the ocular surface^4–7^. When kept in check, Corynebacteria establishes homeostasis with the host and maximizes the host’s ability to resist infection by other pathogens^4,6^; however, disruption of homeostasis can result in disease^5,7^. The mechanisms surrounding Corynebacteria-related homeostasis and disease rely on the ability of the microbe to stimulate IL-17 from γδ T cells in the respective contexts of health and disease. Mycolic acids, a large group of cell wall components shared by Corynebacteria, Mycobacteria, Nocardia and Rhodococcus (phylum Actinomycetota)^8^ have been implicated as adjuvants and/or as inducers of IL-17 responses from γδ T cells^4,7,9–11^. However, resolving the specific microbial factor(s) has remained largely elusive.

A distinguishing feature of Corynebacteria compared to Actinomycetota like *Mycobacterium tuberculosis* is that Corynebacteria lack a type II fatty acid synthase, which results in the surface expression of unique mycolic acids with shorter chain fatty acids, called corynomycolic acids (corynomycolates). The make-up and structure of these corMycs within a single microbial species ranges across saturated, mono-unsaturated, or di-unsaturated acyl tails, and includes sugar conjugates like glucose or trehalose^12^. Notably, in-depth structural and functional characteristics of corMycs in commensal Corynebacteria, as well as the sensing host elements, remain unexplored, limiting our understanding of the roles of corynomycolates in the crosstalk between the microbe and the host, and their potential for clinical development.

In the current study, we report that the lipid fraction of *C. mast* induces IL-17 from γδ T cells *in vitro* and at the ocular surface. Further, using lipidomics and synthetic biology, we identified that trehalose monocorynomycolates (TMCMs) are the primary microbial factors that engage innate and adaptive receptors to stimulate IL-17 from γδ T cells. Mechanistically, we show that TMCMs activate dendritic cells (DCs) by signaling through pattern recognition receptors (PRRs), Toll-like receptor (TLR)-2 and macrophage-inducible C-type lectin (Mincle). Also, TMCMs can activate γδ T cells by engaging the TCR, suggesting that *C. mast* TMCMs are γδ TCR ligands. In addition, TMCMs are sufficient to induce IL-17-mediated immunity *in vivo* and drive ocular surface protection from *P. aeruginosa* infection. Our data highlight TMCMs as promising prophylactic that can bolster IL-17-driven immunity, revealing their potential as an adjuvant in the context of immunization/vaccination and as a potential therapeutic to boost IL-17-driven protective immune responses at barrier sites.

## RESULTS

### Lipid components of *C. mast* induce ocular IL-17A responses

IL-17 from γδ T cells protects the ocular surface from pathogenic infection primarily through the recruitment of neutrophils to the conjunctiva and the production/release of antimicrobial peptides into the tears^4^. Because lipids are common stimulants of γδ T cells^13,14^, we hypothesized that the lipid-rich cell wall and membrane components of *C. mast* harbored the γδ T cell stimulating factor(s). Therefore, we extracted organic-soluble components from *C. mast*, which consisted of hydrophobic, non-protein, and non-nucleic acid molecules (crude lipids). To assess whether *C. mast* lipids could induce IL-17 and proliferation from γδ T cells, we combined crude *C. mast* lipid extracts with dendritic cells (DCs) and bulk γδ T cells from the cervical lymph nodes (LN) of naïve mice and found that *C. mast* crude lipids stimulated IL-17 exclusively from the Vγ4 γδ T cell subset (**Figures 1A and 1B**). Next, crude lipids were applied to the conjunctiva of *C. mast*-naive C57BL/6J wild-type (WT) mice for 7 consecutive days (**Figure 1C**). IL-17A responses were then assessed in eye-draining lymph nodes (DLNs). Compared to the vehicle controls, mice instilled with the crude lipid extract of *C. mast* developed a robust IL-17A response primarily from the Vγ4-positive T cell population, but not the Vγ4-negative population (**Figures 1D and S1**). Instillation of crude *C. mast* lipids also resulted in enhanced recruitment of Vγ4 T cells and neutrophils to the conjunctiva (**Figures 1E and 1F**).

**Figure 1.**
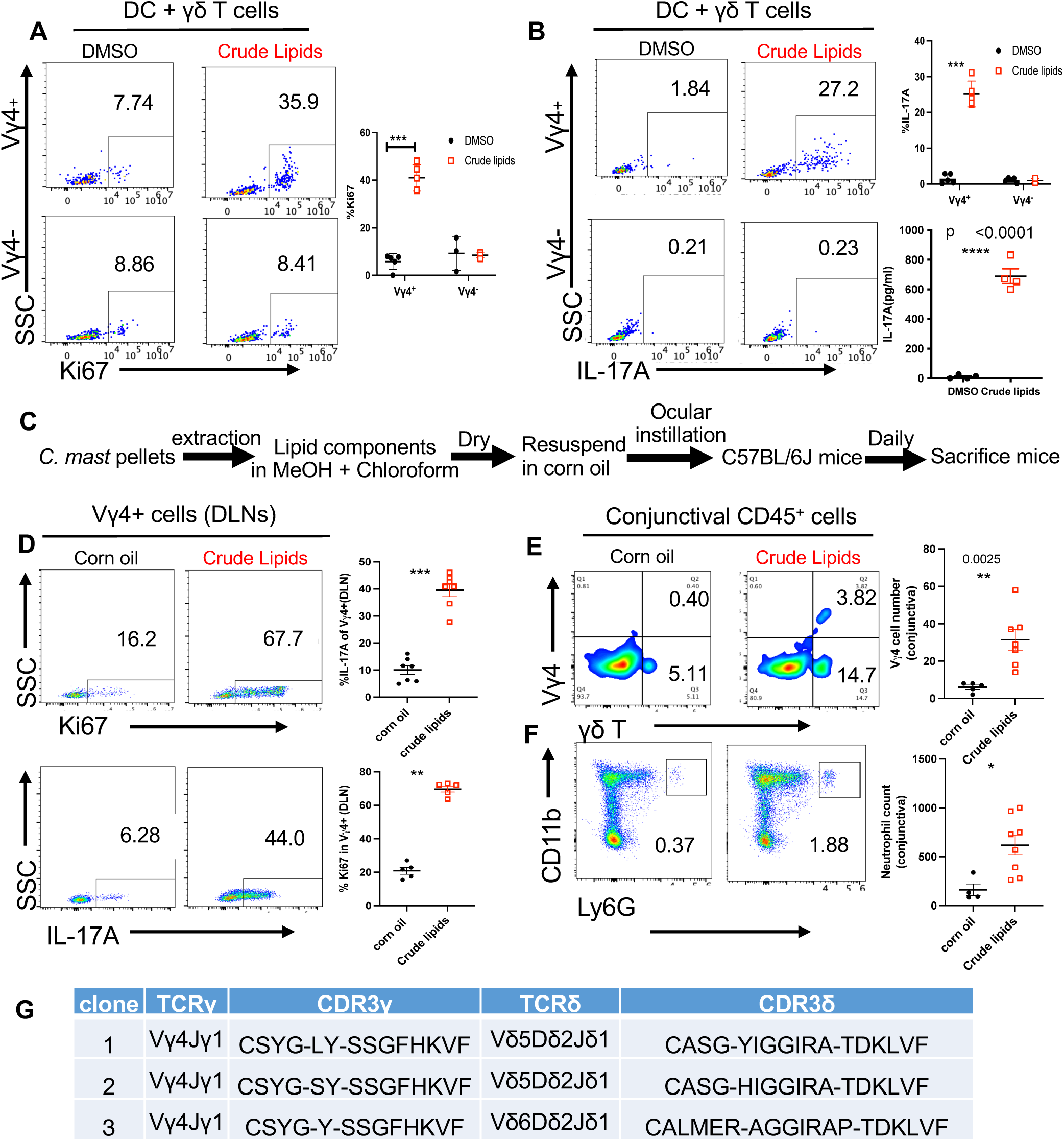
Lipid components of *C. mast* induce ocular IL-17A responses. **(A-B)** Crude lipid components of *C. mast* were extracted by methanol and chloroform and resuspended in DMSO. Sorted γδ T cells were cocultured with CD11c+ DCs in the presence of indicated stimuli for 3 days. Representative FACS plots and dot plots show the percentage of Ki67+ γδ T cells **(A)** or the percentage of IL-17A producing γδ T cells and the amount of IL-17A production in the supernatant measured by ELISA **(B)** when treated with or without lipid components of *C. mast*. Each dot represents one experiment. **(C)** Schematic of experimental design. Crude lipid components of *C. mast* were extracted by methanol (MeOH) and chloroform and resuspended in corn oil. C57BL/6J mice were inoculated with either crude lipid components of *C. mast* (25 µg/per eye) or corn oil daily for 7 days and then sacrificed for profiling immune responses. **(D-E)** Representative flow cytometry plots and dot plots showing the percentage of Ki67+ and IL-17A+ in Vγ4 cells of cervical eye draining LNs (DLNs) **(D)**. Representative flow cytometric plots and total numbers of Vγ4 cells **(E)** and neutrophils **(F)** in the conjunctiva from mice depicted in **(C)**. Each dot represents one animal. (**G**) Vγ4+CD44+CD27- γδ T cells were individually sorted from DLNs of corn oil-associated (single cell number = 131) or crude lipid-associated mice (single cell number =137) and subjected to single-cell TCR sequencing. Animo acid sequences of CDR3 regions of exclusively expanded γδ T cell clones in lipid-associated mice are shown. The results were representative of at least 2 independent experiments (A-B, D-E). Bars represent mean ± SEM with *p<0.05, **p<0.01, ***p <0.001. Statistical significance was determined by the Mann-Whitney test (**A-B, D-F**).

TCR profiling^15^ of CD44^+^CD27^low^ (markers for IL-17A producing γδ T cells) Vγ4^+^ γδ T cells from the DLNs of crude lipid treated mice revealed three clonal populations of γδ T cells that expressed Vγ4Vδ5 or Vγ4Vδ6 TCR chains (**Figure 1G**). Two Vγ4Vδ5 clones utilized germline-encoded Vγ4Jγ1 and Vδ5Dδ2Jδ1 segments without N-region nucleotide additions, whereas the Vγ4Vδ6 clone employed germline-encoded Vγ4Jγ1 segment and Vδ6Dδ2Jδ1 segments with N-region nucleotide insertions. Notably, these TCR clonotypes are utilized by innate-like γδ T cells that acquire the potential of IL-17-production in the thymus^15,16^. Thus, we conclude that *C. mast* lipids contain an activity that can induce an IL-17 response from γδ T cells.

### *C. mast* corynomycolic acids stimulate γδ T cell to produce IL-17

To identify the functional lipid component that stimulates IL-17A production from γδ T cells contained in the crude lipid *C. mast* extract, we first characterized the major lipid components of *C. mast*. Despite reports defining the lipid composition of other Corynebacteria^17^, the lipid composition of *C. mast* was still unknown. Using thin layer chromatography (TLC) and direct injection mass spectrometry analysis, we confirmed the presence of an array of lipids including Phosphatidylglycerol (PG), Cardiolipin (CL), Phosphatidylinositol (PI), free fatty acids (FA) and corynomycolates (corMycs) within the *C. mast* crude lipid extract **(Figures S2A, S2B and S2C)**. In addition, high mass features were detected within the crude lipid extract spectra that were suggestive of conjugated corynomycolates **(Figure S2D)**. Specifically, an acetate adduct series, consisting of putative trehalose monocorynomycolate (TMCM) and trehalose dicorynomycolate (TDCM) species, was identified and further fragmented to resolve acyl-chain content of selected examples from each family (**Figure S2E**). To separate these lipid classes, a hydrophilic interaction chromatography (HILIC) solid phase extraction (SPE1) strategy was adopted to leverage polarity differences in the lipid head groups. Based on polarity, we fractionated the crude *C. mast* lipids into 5 fractions (termed SPE1_F1 to SPE1_F5; **Figure 2A**). Further analysis of these fractions using TLC revealed a successful separation of fatty acids and corynomycolates from phospholipids (PG, CL, and PI) (**Figure 2B**). This separation was further confirmed by direct injection mass spectrometry of the various factions, showing abundant corynomycolate species (between 400-600 mass/charge) in SPE1_F3 and SPE1_F4 (**Figure 2C**).

**Figure 2.**
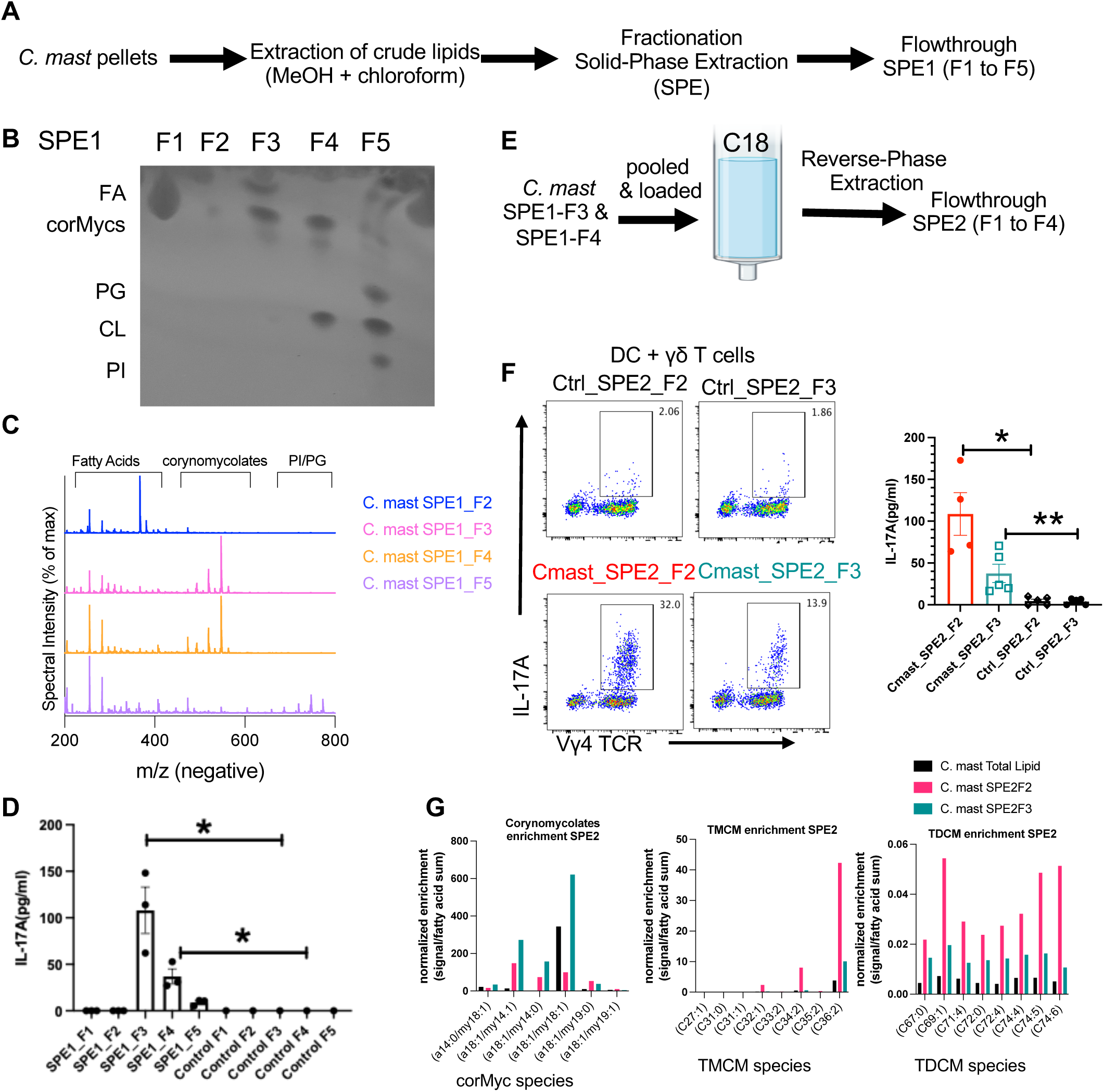
Purified corynomycolate species from *C. mast* can stimulate IL-17A production *in vitro*. **(A-C)** Experimental design to fractionate *C. mast* lipidome using solid phase extraction **(A)** and analyzed by thin layer chromatography(TLC) analysis **(B)**: fatty acids (FA), corynomycolates (corMycs), phosphatidylglycerol (PG), cardiolipin(CL), phosphatidylinositol (PI). (**C**) Lipid fractions from *C. mast* lipidome were examined by liquid chromatography-mass spectrometry (LC-MS). F1 showed a redundant pattern to F2 and is not shown to preserve space. (D) Each lipid fraction of *C. mast* was added in cocultures with sorted γδT cells and CD11c+ DCs for 3 days; the amount of supernatant IL-17A was measured by ELISA. Each dot represents one experiment. (E) Experimental design to separate corynomycolates. (F) Sorted γδT cells were cocultured with CD11c+ DCs in the presence of SPE2_F2 or SPE2_F3 of *C. mast* for 3 days. Representative flow cytometry data and bar plots showing the percentage of IL-17A+ γδ T cells (**F, left**) and the amount of IL-17A secreted in the supernatant (**F, right).** Each dot represents one experiment. (G) The amount of major corynomycolate, TMCM, and TDCM species in total lipids, SPE2_F2 and SPE2_F3 are displayed. Signals were normalized to the sum of free fatty acid signals in their own samples as a surrogate for enrichment. All signals were measured in negative mode ionization with free fatty acids using the neutral loss of 44 series, corynomycolates via the major α-chain product ion, and both TMCM and TDCM via the conserved 323 trehalose-associated product ion. Bars represent mean ± SEM with *p<0.05, **p<0.01, ***p <0.001. Statistical significance was determined by Mann-Whitney test (**D, F**).

To assess the relative ability of each lipid fraction to stimulate IL-17A production, both the lipid fraction and their respective mock extracts (from mock loaded SPE columns) were added to *in vitro* co-cultures of DCs and γδ T cells. The IL-17 response was analyzed by measuring the percentage of IL-17A^+^ γδ T cells and the amount of IL-17A released into the supernatant. SPE1_F1 and F2 fractions and their corresponding mock extracts did not induce IL-17A production by γδ T cells. SPE1_F3 fraction containing fatty acids and corynomycolates induced the highest level of IL-17A production from γδ T cells, followed by SPE1_ F4 fraction containing corynomycolates and cardiolipin. The phospholipid-containing SPE1_F5 fraction had only a minimal IL-17A stimulatory effect (**Figures 2D and S3**).

To further purify the active factor from the mixture of corynomycolates and free fatty acids in SPE1_F3 and F4, we subsequently performed reverse phase SPE separation (referred to as SPE2) on a mixture of the active fractions from SPE1 (**Figure 2E**). We then used the fractions of the SPE2 separation to stimulate co-cultures of γδ T cells and DCs, and found that SPE2_fraction 2 (SPE2_F2) and SPE2_fraction 3 (SPE2_F3) both stimulated IL-17 production (**Figure 2F**). To identify the specific lipid components responsible for IL-17 production, a custom targeted liquid chromatography-tandem mass spectrometry (LC-MS/MS) method was used on fractions SPE2_F2 and SPE2_F3 to specifically profile levels of corynomycolates, TMCMs, TDCMs, and free fatty acids. Fraction SPE2_F2 was enriched for sugar conjugated forms of corynomycolates like TMCM and TDCM, while fraction SPE2_F3 was enriched for corynomycolates without sugar conjugation (**Figure 2G**). While these results implicated naked or sugar-conjugated forms of corynomycolates or similar lipids as the primary driver of IL-17A stimulation by *C. mast*, the relative contribution of the different corynomycolate conjugation forms responsible for this activity could not be resolved using this approach.

### Trehalose monocorynomycolate (TMCM) induces IL-17A production in γδ T cells

To identify the specific corynomycolates with IL-17A stimulatory capacity, we synthesized a representative naked (unconjugated) corynomycolate (corMyc (a18:1/my18:1)), a trehalose monocorynomycolate (TMCM (a18:1/my18:1)) and a trehalose dicorynomycolate (TDCM (a18:1/my18:1, a18:1/my18:1)) based on the most abundant corynomycolate species detected in the crude lipid extract from *C. mast* (**Figures 3A** and **2G**). The IL-17A stimulatory capacity of these compounds was again evaluated in DC and γδ T cell co-cultures. While TMCM(a18:1/my18:1) (a.k.a., TMCM(36:2)) potently stimulated IL-17A production, neither corMyc (a18:1/my18:1) (a.k.a., corMyc(36:2)) nor TDCM(a18:1/my18:1, a18:1/my18:1)(a.k.a., TDCM(72:4)), showed significant IL-17A inducing effects (**Figure 3B**). Taken together, we identified TMCM as the primary *C. mast*-derived stimulant that activated Vγ4^+^ γδ T cells.

**Figure 3.**
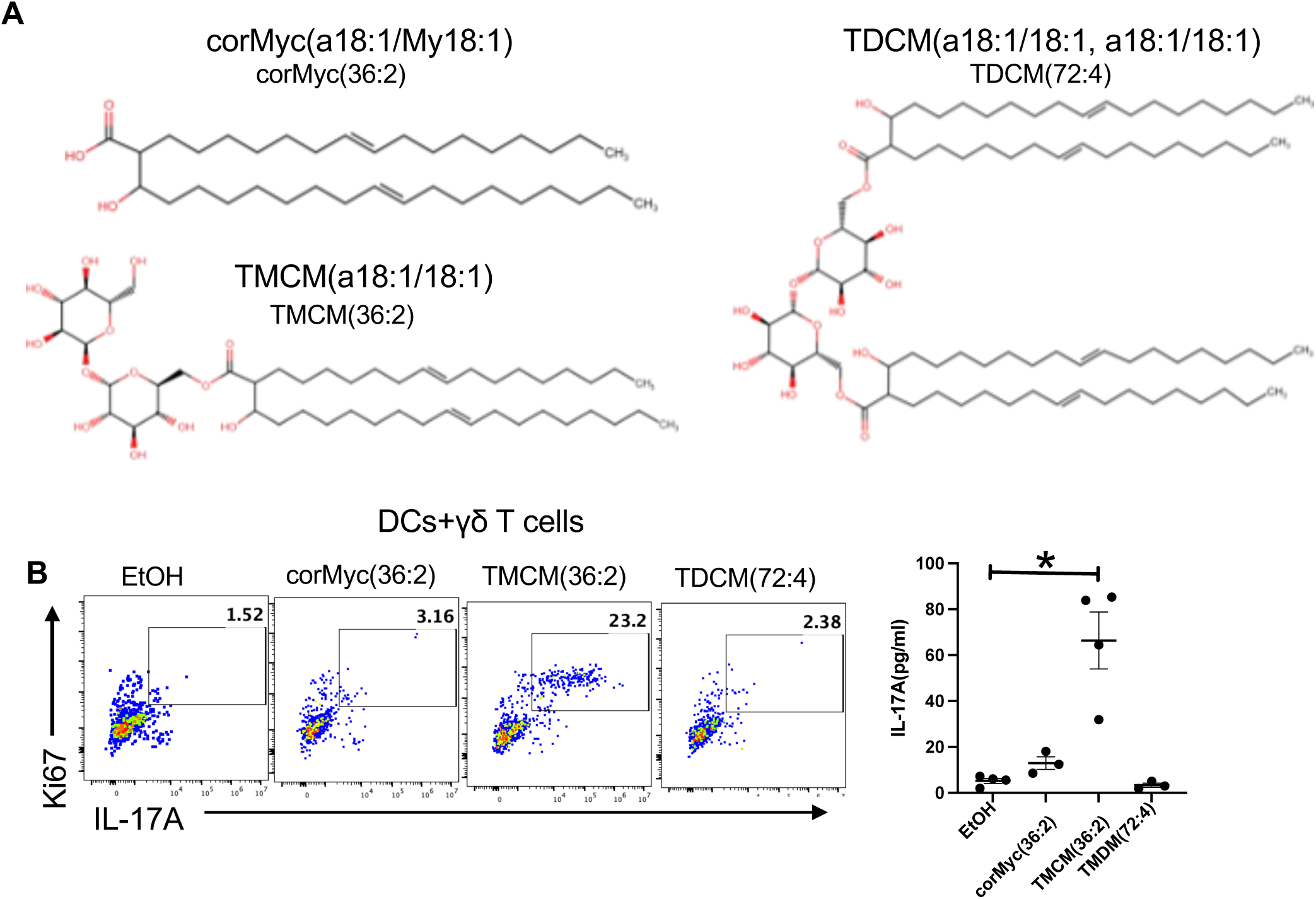
Trehalose monocorynomycolate (TMCM), but not trehalose dicorynomycolate (TDCM) or unconjugated corynomycolate, is a potent IL-17A stimulant. (A) Schematic of synthetic corMyc(a18:1/My18:1) (corMyc(36:2)), D-(+)-trehalose 6- monoMyc(a18:1/My18:1) (TMCM(36:2)) and D-(+)-trehalose 6,6’-diMyc(a18:1/My18:1) (TDCM(72:4)). (B) Synthetic corMyc(36:2), TMCM(36:2), or TDCM(72:4) were dissolved in ethanol (EtOH). 1 µg/ml of each synthetic lipid was added in cocultures with sorted γδT cells and CD11c+ DCs for 3 days. The frequency of Ki67+ IL-17A producing Vγ4+ T cells and the level of IL-17A in the culture supernatant are shown. Each dot represents one experiment. Bars represent mean ± SE. *p<0.05, **p<0.01, ***p <0.001. Statistical significance was determined by Mann-Whitney test (B).

### Recognition of TMCM(36:2) by TLR2 and Mincle contributes to induction of IL-17A from **γδ T cells**

After identifying the specific stimulant, we used the synthetic TMCM to dissect the downstream signaling events leading to IL-17 production. DCs are essential for the induction of IL-17A from γδ T cells after stimulation with TMCM(36:2) (**Figure S4A**). We therefore examined how DCs may sense and respond to TMCM(36:2). We first analyzed the cytokine secretion profile of DCs stimulated with TMCM(36:2), and showed that TMCM(36:2) induces the production of IL-1α, TNF-α, MCP-1, IL-1β and IL-6 by DCs (**Figure 4A**) while IL-23, IFN-β, ΙL-10, IL-27, IL-12p70, GM-CSF were below the limit of detection (data not shown). Since TLR2 and Mincle can recognize glycolipids from Corynebacteria^18^, we examined whether TLR2 and/or Mincle on DCs was required for TMCM(36:2) induction of IL-17. TLR2 deficient DCs exhibited a marked reduction (p<0.05) in the production of IL-1α, TNF-α, IL-1β, and IL-6 (**Figure 4B**). Anti-Mincle pretreated DCs displayed reduced IL-1α and TNF-α in response to TMCM (36:2) stimulation, compared to IgG-pretreated DCs (**Figure 4C**). In line with this, γδ T cells cocultured with TMCM(36:2)-pulsed TLR2-deficient or Mincle-blocked DCs produced less IL-17A (**Figure 4D-E**), suggesting that recognition of TMCM(36:2) by TLR2 and/or Mincle on DCs influences IL-17A production by γδ T cells though cytokines IL-1, IL-6 and/or TNF-α. Blockade of IL-1R signaling almost completely abolished IL-17A production in response to synthetic TMCM(36:2) (**Figure 4F**). To investigate the roles of IL-6 and TNF-α in IL-17A production, we treated cocultures with individual blocking antibodies or a combination of both. Blocking IL-6 or TNF-α individually did not reduce IL-17 production; however, simultaneous blockade of IL-6 and TNF-α led to a partial decrease in IL-17 production (**Figures S4B and 4G**). These findings demonstrate that TMCM(36:2) activates the innate immune system at least in part through TLR2 and/or Mincle receptors, triggering an IL-1-dependent pathway similar to wild-type *C. mast* stimulation of IL-17 production from γδ T cells^4^.

**Figure 4.**
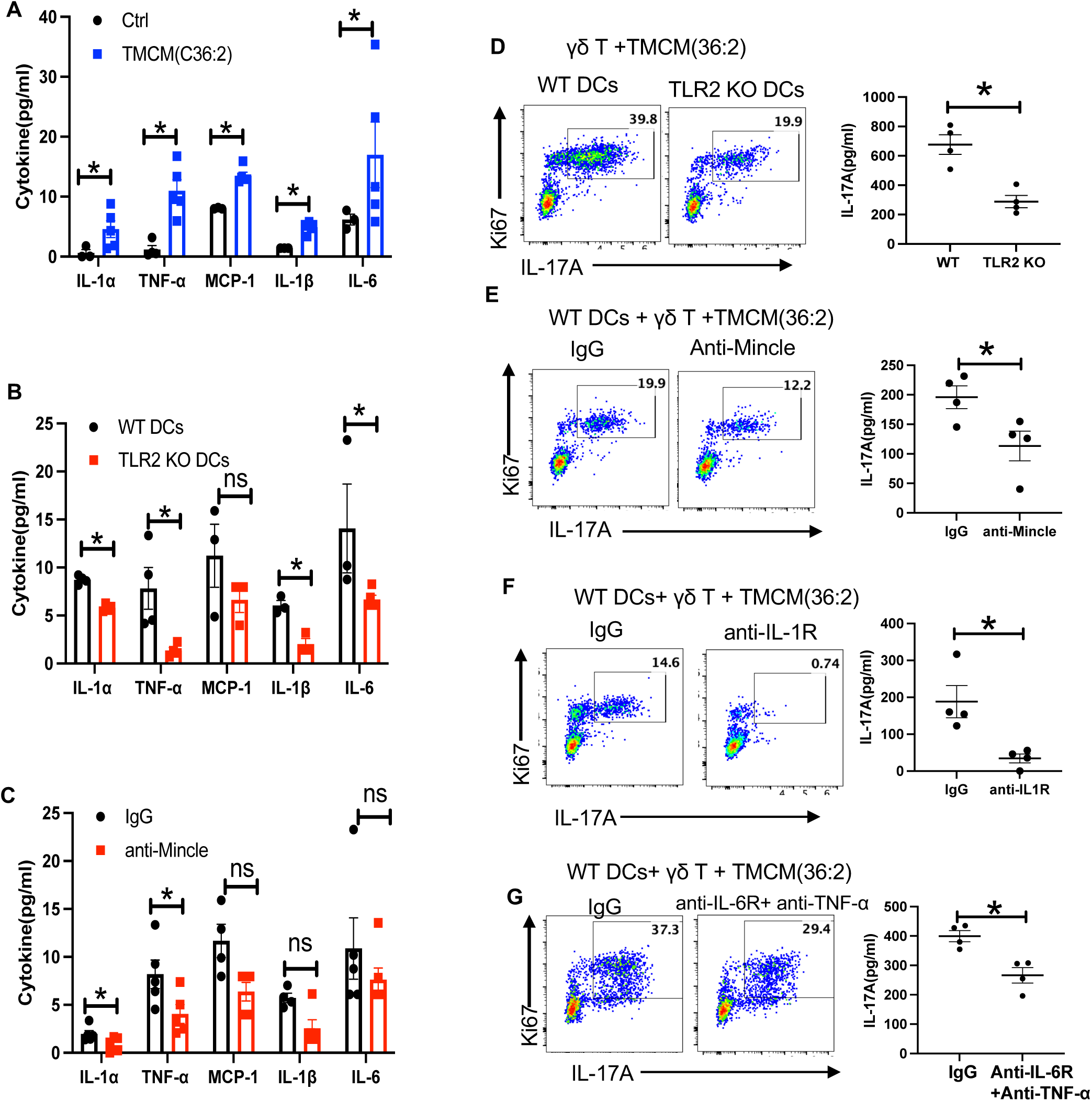
TLR2 and Mincle expressed by DCs sense TMCM and regulate γδ T cell IL-17A production. (**A–C**) DCs were stimulated with either vehicle control ethanol or 1 µg/ml synthetic TMCM(36:2) for 2 days. (**A**) WT DCs, (**B**) TLR2 deficient DCs, or (**C**) WT DCs pretreated with IgG or anti-Mincle antibody, were stimulated with TMCM(36:2) for 2 days. Cytokine levels in the culture supernatants were measured by a Legendplex mouse inflammation panel for 13 analytes. Bar plots list five cytokines that are induced by TMCM. Each dot represents one experiment. (**D**) Sorted γδ T cells were cocultured with WT or TLR2 deficient CD11c+ DCs in the presence of 1 µg/ml synthetic TMCM(36:2) for 72 hours. Representative FACS plot showing the percentage of Ki67+ IL-17A+ γδ T cells and the scatter plot showing the level of IL-17A in the culture supernatant. Each dot represents one experiment. (**E-G**) FACS plots showing the percentage of Ki67+ IL-17A+ γδ T cells and the scatter plot showing levels of IL-17A production in the supernatants from γδ T cells co-cultured with WT DCs (**E**) with or without anti-Mincle for 48 hours; or (**F)** with or without anti-IL-1R for 48 hours; or (**G)** cultures with or without anti-IL-6R + anti-TNF-α for 48 hours. Each dot represents one experiment. Bars are mean ± SEM. *p<0.05, **p<0.01, ***p<0.001. Statistical significance was determined by the Mann-Whitney test (A-G).

### TCR signaling is critical for IL-1R1 upregulation to support IL-17A response of γδ T cells responding to TMCM

Even though TMCM-mediated IL-17 responses from γδ T cells appeared to rely on innate signaling in DCs, and the conserved γδ TCR repertoire *in vivo*, it was still uncertain whether TCR signaling was required for γδ T cell IL-17 production in response to TMCM. To address this, we used ‘knock-in CD3ζ switch’ mice (CD3ζ 6Y/6Y(6F(induced)))^19^, in which tyrosine (Y) in immunoreceptor tyrosine-based activation motif (ITAM) of CD3ζ proteins was mutated to phenylalanine (F) after tamoxifen induction (= mutant CD3 mice), impairing γδ TCR signaling (**Figures S5A and S5B**)^19^. Eight days after tamoxifen induction, γδ T cells were isolated from peripheral LNs and analyzed for cell surface expression of CD44+ CD27^−^ (marker for IL-17A producing γδ T cells) and IL-17 production after PMA/Ionomycin stimulation. By both criteria, γδ T cells with IL-17A-producing phenotype in CD3 mutant mice were unchanged (**Figures 5A and 5B**), indicating that γδ T cell development and effector function remained unaffected. Importantly, IL-1R1 expression by CD44+ CD27-γδ T cells was reduced in CD3 mutant mice (**Figure 5C**) and their IL-1R signaling after stimulation with IL-1β *in vitro* was decreased, as indicated by pErk and pP38 levels (**Figures 5D,5E**). Of note, only IL-1R1 expression was affected, whereas IL-7Rα expression remained normal (**Figure S5C**). In line with their decreased IL-1R signaling, mutant CD3 γδ T cells exhibited dramatically decreased proliferation and IL-17A production in response to synthetic TMCM-pulsed DCs (36:2) (**Figure 5F**). In view of the nonredundant role of IL-1R signaling in γδ T cells for IL-17A production (**Figure 4F**), we propose that TCR signals are essential for IL-17A production of Vγ4 T cells in response to TMCM, at least in part by controlling their intrinsic IL-1R1 expression.

**Figure 5.**
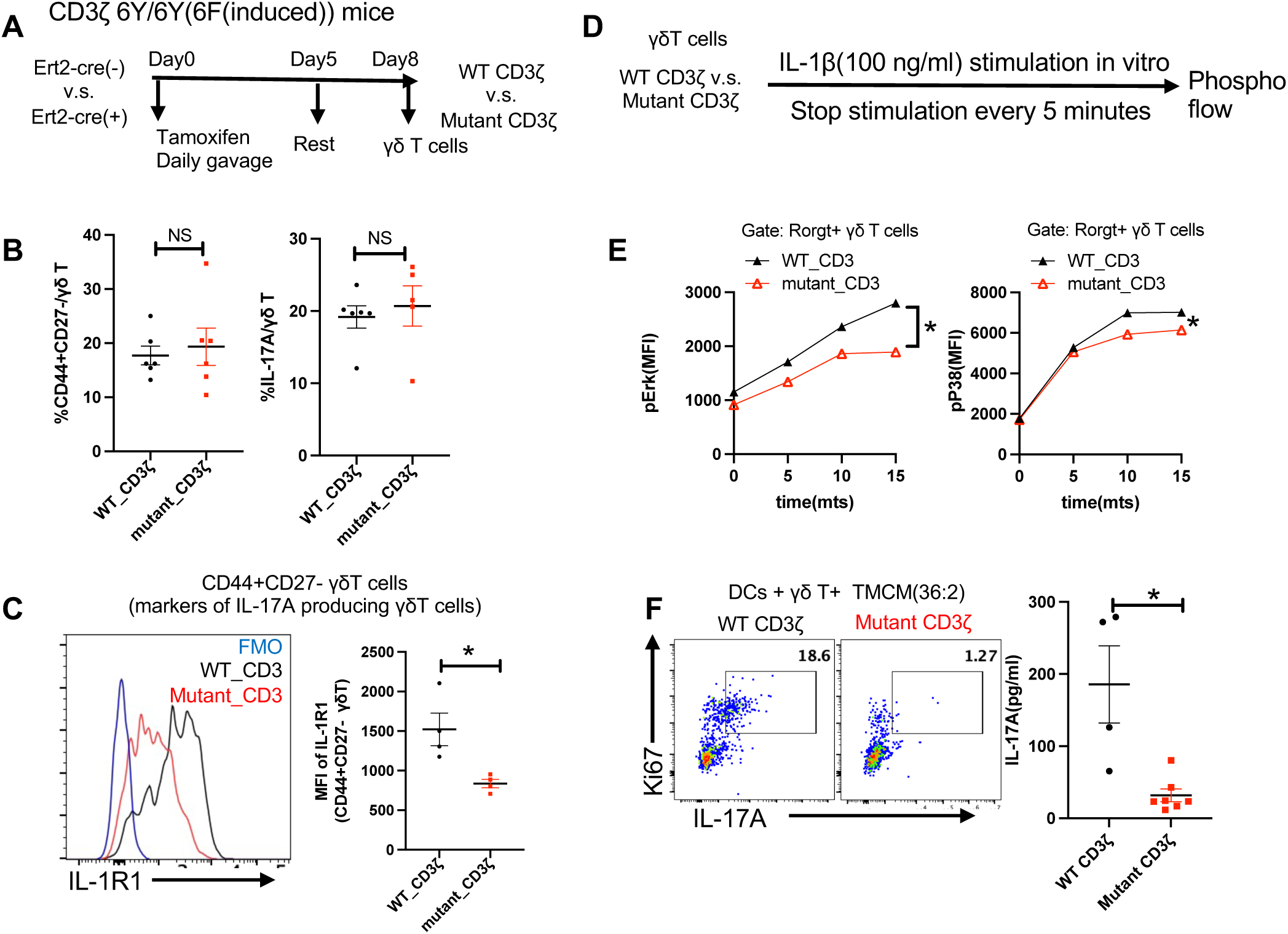
γδ TCR signaling supports IL-1R1 expression and is required for IL-17A production in response to TMCM. (A) Experimental design for inducing expression of WT or mutant CD3ζ in Ert2-cre(-) CD3ζ 6Y/6Y(6F(induced)) (WT CD3ζ) and Ert2-cre(+) CD3ζ 6Y/6Y(6F(induced)) (Mutant CD3ζ) mice. (B) Percent of CD44+CD27- (left) and IL-17A+ (right) cells among γδ T cells in LNs of WT and mutated CD3ζ mice 8 days after tamoxifen induction. Each dot represents an individual mouse. (C) Overlaid histograms and scatter plots show the expression of IL-1R1 on CD44+CD27- γδ T cells from WT and mutated CD3ζ mice 8 days after tamoxifen treatment. Each dot represents an individual mouse. Fluorescence minus one (FMO) as controls. **(D-E)** γδ T lymphocytes purified from WT and mutated CD3ζ mice on day 8 after tamoxifen treatment were stimulated with IL-1β and analyzed by phospho-flow (**D**). The mean fluorescence intensity (MFI) of phospho-Erk (left) and phospho-P38 (right) in Rorgt+ γδ T cells (γδ T cells with IL-17A producing phenotype) at indicated time points after IL-1β stimulation is displayed. Representative data from three independent experiments are shown (**E**). **(F**) γδ T cells were sorted from WT or mutated CD3ζ mice 8 days after tamoxifen treatment and co-cultured with DCs in the presence of TMCM(36:2) for 48 hours. A representative FACS plot illustrates the proportion of Ki67+ IL-17A+ γδ T cells in WT or mutant CD3ζ γδ T cells. The scatter plot displays the IL-17A levels in culture supernatants quantified by ELISA. Data are pooled from two independent experiments. The results were representative of at least 2 independent experiments. Bars represent mean ± SEM with *p<0.05, **p<0.01, ***p<0.001.Statistical significance was determined by Mann-Whitney test (**B-C, F)** and two-way ANOVA (**E**).

### Corynomycolate-deficient *C. mast* mutant fails to stimulate protective IL-17 response from γδ T cells

To show that corynomycolates are necessary for *C. mast*’s colonization and induction of protective immunity at the ocular surface, we screened our previously described *C. mast* transposon (Tn) mutant library for biofilm formation, a phenotype potentially affected by mycolic acid disruption^20,21^. We identified a mutant that has its Tn insertion in the *C. mast* FadD32 Polyketide Synthase gene (termed *Pks/FadD32::Tn*) **(Figure S6A)**, which is essential for the production of mycolic acids in Mycobacteria and Corynebacteria^22^. To verify that this gene is essential for *C. mast* corynomycolic acid production, we analyzed its lipidome and found that *Pks/FadD32::Tn* completely lacked corynomycolic acids (**Figures 6A, S6B and S6C**). This mutant is also unable to colonize the eye, despite developing a strong biofilm *in vitro*, presumably because its disrupted mycomembrane cannot protect *C. mast* in the antibacterial environment of the ocular surface (**Figures 6B and 6C**). We can conclude from this that corynomycolic acids provide bacterial structure in the context of biofilm formation and contribute to the ability of *C. mast* to survive the hostile ocular environment. There are likely other mechanisms that contribute to colonization given that *C. glutamicum* is unable to colonize the eye^4^, although it contains similar corynomycolates as *C. mast*^12^ (**Figure S6D**).

**Figure 6.**
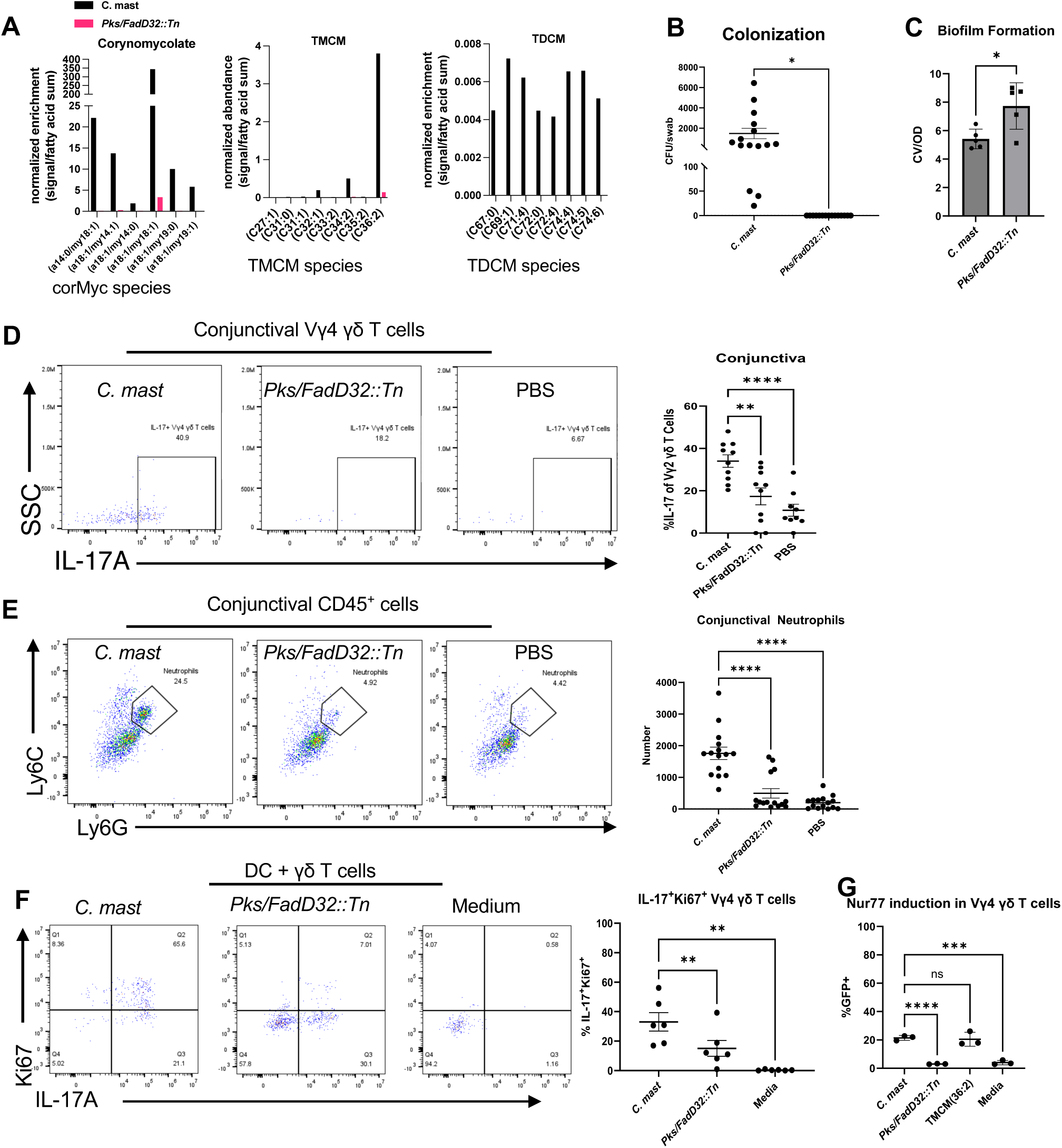
Corynomycolate-deficient *C. mast* Tn mutant fails to stimulate protective IL-17 response. **(A)** Analysis of corynomycolate profiles of *C. mast* or mutant *Pks/FadD32::Tn* was performed using LCMS. Comparison of relative enrichment was performed between *C. mast* and *Pks/FadD32::Tn* by normalizing signals to the sum of free fatty acid signals. **(B)** Three-week-old C57BL/6 mice were inoculated every other day for a total of three inoculations with 5x10^5^ CFU/eye of WT *C. mast* or the corynomycolate deficient *C. mast* mutant *Pks/FadD32::Tn*. One week post final inoculation, eyes were swabbed and streaked on LBT plates with Fosfomycin. Graphs show the number of CFU/swab, where symbols represent individual mice from three independent experiments. Significance was determined using Welch’s T test (*p=0.0115). **(C)** *C. mast* or mutant *Pks/FadD32::Tn* were grown in 48-well tissue culture plates for 48 hours. Bacterial growth was quantified at OD600 (OD), wells were washed, and the biofilm on the plate surface was stained with 0.1% crystal violet and quantified at OD550 (CV). Graph shows the amount of biofilm normalized to growth (CV/OD), where symbols represent individual experiments. Significance was determined by Welch’s T test (*p=0.0304) (**D-E**)Two weeks after the final inoculation (described in **B**), mice were sacrificed, and conjunctivas were harvested. Single cells suspensions were stimulated with PMA/Ionomycin and analyzed for IL-17+ Vγ4 γδ T cells in the conjunctiva (**D**); and assessed for number of neutrophils in the conjunctiva by flow cytometry **(E)**. For all graphs, symbols represent data for individual mice from two **(D)** or three **(E)** independent experiments. Bars represent average ± SEM. Differences were determined to be non-significant using an ordinary one-way ANOVA (**p=.0036, ***p=.0001, ****p<0.0001). **(F-G)** 1x10^4^ γδ T cells were obtained from WT mice (**F**) or Nur77^GFP^ reporter mice(**G**) which pre-associated with *C. mast* and cocultured with 1x10^5^ DCs pretreated with 1x10^5^ CFU of either WT *C. mast* or *Pks/FadD32::Tn* 24 hours or treated with TMCM(36:2). After 72 hours of coculture, γδ T cells were assessed for percent of IL-17+Ki67+(**F**) and Nur77^GFP^(**G**) in Vγ4 γδ T cells by flow. Bars are mean ± SEM. Symbols represent individual experiments. Significance was determined using a repeated measure one-way ANOVA (**p=.0006).

Since corynomycolic acids in *C. mast* are necessary for induction of IL-17 from γδ T cells (**Figure 3**) we hypothesized that *Pks/FadD32::Tn* would be unable to induce IL-17 from γδ T cells. Indeed, application of *Pks/FadD32::Tn* to the ocular surface did not recruit IL-17 producing γδ T cells, and did not attract neutrophils to the conjunctiva (**Figures 6D and 6E**). Because this lack of *in vivo* IL-17 by *Pks/FadD32::Tn* could be due to its lack of ability to colonize, we examined whether the mutant stimulated IL-17 from γδ T cells *in vitro.* In a coculture system of DCs and γδ T cells, *Pks/FadD32::Tn* failed to stimulate IL-17 production compared to wild-type *C. mast*, confirming directly that corynomycolates are responsible for the ability of *C. mast* to induce IL-17 from γδ T cells (**Figure 6F**). To identify whether inability of the mutant to induce IL-17 was due to a lack of recognition by innate immune cells, we exposed bone marrow derived DCs to WT vs mutant *C. mast* to look for possible differences in maturation, as assessed by the markers CD80 and CD86. Expression of the co-stimulatory marker, CD80, was slightly reduced, whereas CD86 expression did not differ between the *Pks/FadD32::Tn* and WT *C. mast* (**Figure S7A**). Of note, DCs produced significantly less IL-1β when stimulated with the *Pks/FadD32::Tn* mutant compared to WT *C. mast* (**Figure S7B**). These data support the notion that corynomycolic acids are recognized by innate immune cells.

To assess whether corMycs in *C. mast* are necessary to induce TCR signaling, we cocultured *Pks/FadD32::Tn* with DCs and Nur77^GFP^ γδ T cells and found that, unlike WT *C. mast*, *Pks/FadD32::Tn* did not induce Nur77 signaling. In contrast, TMCM(36:2) induced Nur77 expression similarly to WT *C. mast.* This highlights the role of corMycs, specifically TMCM(36:2) (**Figure 6G**), in TCR signaling. Therefore, we propose that corMycs are necessary and sufficient for *C. mast* colonization and for immune recognition by γδ T cells.

### Synthetic TMCM induces *in vivo* immunity and protects the ocular surface from bacterial keratitis

Since corynomycolates are essential for *in vivo* immunity (**Figures 6D and 6E**), we examined whether synthetic TMCM (36:2) was sufficient to induce protective immunity at the ocular surface. Ocular instillation of synthetic TMCM(36:2) triggered a robust IL-17 response in Vγ4 T cells of the eye DLN (**Figure 7A-B**) along with recruitment of Vγ4^+^ T cells and neutrophils to the conjunctiva (**Figure 7C**), without affecting corneal integrity (**Figure 7D**), whereas the unconjugated corMyc (36:2) or TDCM(72:4) failed to stimulate *in vivo* immunity (**Figure S8A-D**). Lastly, we asked if TMCM(36:2) treated mice would be protected from ocular surface challenge with *Pseudomonas aeruginosa,* similarly to *C. mast* colonized mice^4^. Mice treated with TMCM had significantly fewer *P. aeruginosa* CFU isolated from the eye, similarly to mice inoculated with whole *C. mast*, whereas mice treated with unconjugated corMyc or with the mutant *Pks/FadD32::Tn* had a similar *P. aeruginosa* burden as control mice receiving PBS (**Figure 7E**). These data show that TMCM(36:2) is sufficient to protect the ocular surface from pathogenic challenge and highlight the therapeutic potential of TMCM (36:2) as a novel strategy to combat ocular infections.

**Figure 7.**
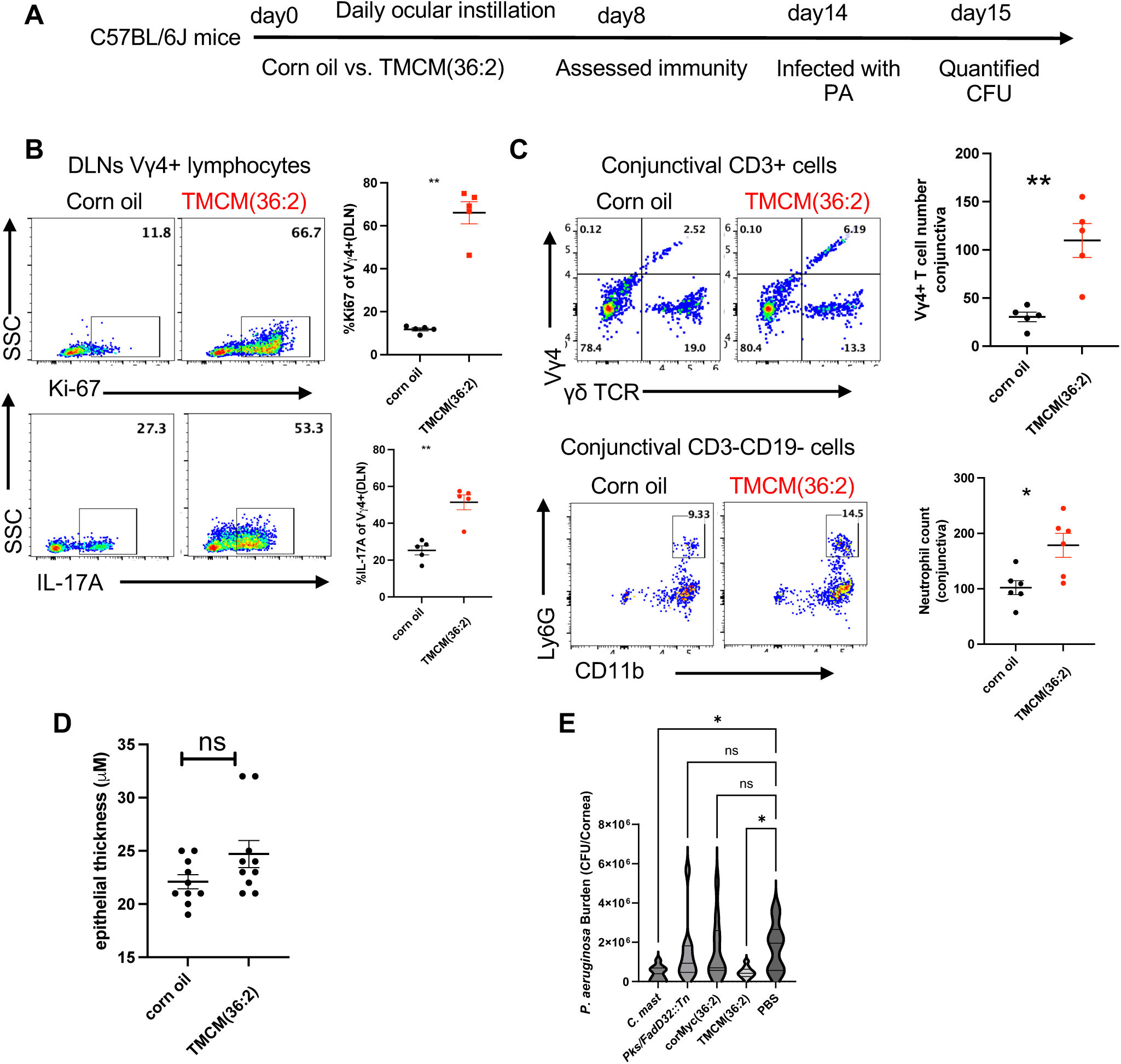
Synthetic TMCM elicits *in vivo* immunity and protects the ocular surface from bacterial keratitis. (**A**) Experimental design to assess ocular immune responses to synthetic TMCM(36:2). (**B-C**) Corn oil (vehicle control) or 25 µg synthetic TMCM(36:2) was instilled to conjunctivas on C57BL/6J mice daily for 7 days. **(B)** Representative flow cytometric analysis and dot plots show the percentage of Ki67+ or IL-17A+ Vγ4 cells of eye-draining cervical LNs (DLNs) with PMA and ionomycin stimulation. **(C)** Representative flow cytometric analysis and total numbers of Vγ4 cells and neutrophils infiltrating the conjunctiva of mice. Each dot represents one animal. Representative data for two experiments. Statistical significance was determined by Mann-Whitney test (**B-C**). (D) The corneal epithelial thickness of mice described in (B) was acquired by anterior segment optical coherence tomography (OCT) and plotted. Each dot represents one animal. Data are pooled from two independent experiments. (E) Four-week-old C57BL/6 mice were either inoculated every other day for a total of three inoculations with 5x10^5^ CFU/eye of WT C. mast or *Pks/FadD32::Tn*, or every day for two weeks with 25 μg of synthetic TMCM(36:2) or corMyc(36:2). 24 hours after the final lipid inoculation, mice were challenged with 1x10^5^ CFU/eye of *P. aeruginosa.* The next day, whole eyes were obtained, homogenized in PBS, and serially diluted on LB agar. (n=10 mice per group) Significance was determined using standard one-way ANOVA (*p=.0202). The results were representative of at least 2 independent experiments(B-C,E). Bars represent mean ± SEM. *p<0.05, **p<0.01, ***p <0.001.

## DISCUSSION

Our earlier studies identified a causal relationship between *C. mast*, an eye-colonizing microbe, and ocular surface health and disease^4^. Colonization of *C. mast* on the ocular surface induces an immune response mediated by IL-17 production from γδ T cells that protects the eye from bacterial and fungal corneal infection. In the current study, we demonstrate that corMycs are essential for both *C. mast* ocular surface colonization and induction of protective immunity. We uncover that TMCM, but not TDCM, is the primary immunostimulatory component of *C. mast,* and is both necessary and sufficient to stimulate protective IL-17 production from γδ T cells. Mechanistically, the innate receptors TLR2 and Mincle on DCs, and the engagement of the γδ TCR, are required for the full induction of the γδ T cell IL-17 response by TMCM.

### The ability to persist on the ocular surface is a prerequisite for successful commensalism and depends on corynomycolates

Corynebacteria have a mycomembrane which is the outer part of the cell wall complex and contributes to virulence, colony morphology, colonization of the host, and cell permeability^23,24^. It is a multi-component structure whose backbone is composed of corMycs. The generation of a Tn mutant library of *C. mast* yielded a mutant lacking corMycs that could not colonize the ocular surface, confirming the role of corMycs in colonizing the eye. The unique role of corMycs in *C. mast* colonization and immune activation is underscored by *C. glut*, which, despite possessing corynomycolates, fails to colonize and elicit an immune response at the ocular surface. A potential explanation for the loss of colonization in *Pks/FadD32::Tn* is due to a disruption of the mycomembrane, which could expose *Pks/FadD32::Tn* to hostile factors present on the ocular surface, such as lysozyme. Additionally, disruption of the mycomembrane of *Pks/FadD32::Tn* may have compromised its ability to act as a lipid anchor for adhesion molecules that bind to the mucosal surface. Although our Tn mutant exhibited more biofilm formation *in vitro,* it appears that the ability to make a biofilm may be necessary, but is insufficient, for successful ocular surface colonization.

### The gaps in knowledge that are filled by our study

The TMCM-reactive subset of IL-17A-producing γδ T cells fit the definition of innate-like γδ T cells, which are pre-programed during thymic development to rapidly produce IL-17A in response to cytokine stimuli in the periphery^25–27^. Several studies established the essential role of γδ TCR signaling in thymic development and lineage specification of γδ T cells, including the induction and expression of IL-7R and IL-1R^26–28^. However, the role of TCR signaling in their IL-17 response after exiting the thymus to the periphery is controversial, due to (a) lack of inducible TCR signaling pathway mutants, which precluded dissociation of effects on development from effector function in the periphery^28–30^; (b) paucity of known γδ TCR ligands^31^ and (c) limited data on γδ TCR and innate receptor crosstalk in γδ T cells.

Our data help address these gaps, in that:

a. we used a conditional mutant of CD3ζ, which impairs TCR signaling in mature extra-thymic γδ T cells^19^. Reduced IL-17 production by the mutant γδ T cells in response to TMCM demonstrates that TCR signaling is required for IL-17A production by these innate-like γδ T cells in the post-thymic stage. This conclusion is supported by Nur77 upregulation by TMCM, indicating TCR engagement. Additionally, the absence of corynomycolates in the corMyc deficient *C. mast* mutant, *Pks/FadD32::Tn*, leads to a reduction in TCR signaling,
b. our data identify TMCM as a novel and biologically relevant ligand for at least three expanded γδ TCRs that function in IL-17-committed γδ T cells.
c. Cytokines produced by myeloid cells (in our model, DC responding to TMCM through TLR2 and Mincle) include production of IL-1 which binds to the IL-1R on γδ T cells and is necessary and nonredundant for their activation to produce IL-17 (**Figure 4F** and ref.^4^). Notably, we show that γδ TCR signaling supports endogenous IL-1R1 expression in γδ T cells. This has not been reported previously and may have important physiological relevance. Namely, although it has been reported that γδ T cells can produce IL-17 in response to external cytokine cues without the need for γδ TCR engagement, this was observed in culture, likely in the presence of supra-physiological amounts of exogenous cytokines^30,32^. Our data emphasize that, under physiological conditions, γδ TCR ligation may be necessary to potentiate this response to biologically significant levels.

### Translational relevance

Infectious keratitis occurs frequently in contact lens wearers, and can be associated with altered commensal communities^33^. Our lab and others demonstrated that the topical association of an ocular commensal, *Corynebacterium mastitidis,* or coagulase-negative Staphylococci, can protect mice against keratitis induced by *Pseudomonas aeruginosa*^4,34^, raising the possibility of using probiotics as therapy for keratitis. However, the challenges to using live commensals are its uncontrolled overgrowth in immune-compromised or excessive inflammation in hyper-reactive individuals^5^, which can damage corneal function. Our approach demonstrates that microbiome-informed therapies can be made safer and more versatile, by demonstrating that a highly purified synthetic form of TMCM successfully substitutes for the whole commensal in protection from P. aeruginosa infection. This approach enables clinical-grade production and precise therapeutic calibration, paving the way for clinical application.

### Limitations of the study

Although our data show that γδ TCR engagement by TMCM plays a key role in activating IL-17 production from innate-like Vγ4+ T cells, the possible contribution of other *C. mast*-sensing molecules on these γδ T cells have not been investigated in this study. This issue is partly addressed in a separate study, where we investigate the function and mechanism of intrinsic TLR2 expression on responses of γδ T cells to *C. mast*^35^.

It has been reported that γδ TCRs can bind antigens directly, without dependence on classical antigen-presenting molecules. Our current data do not address whether TMCM binds and is recognized directly by the γδ TCR, or whether it requires presentation by DC. Studies are ongoing to determine the mode of recognition of the TMCM antigen.

### In conclusion

Our study identifies TMCM as the primary mediator of commensal-driven protective immunity at the ocular surface. These corMycs are necessary for colonization of the eye, and mediate interactions with innate and adaptive immune cells. Specifically, *C. mast* TMCM is necessary and sufficient for *C. mast* driven IL-17 production from γδ T cells. This process involves an interaction with the innate receptors TLR2 and Mincle and production of IL-1 by DC, as well as engagement and activation of the γδ TCR, whose signaling additionally supports endogenous expression of IL-1R1. We conclude that TMCM mimics the ability of the whole commensal to protect the eye from infection by stimulating local protective immunity. Our findings provide a rationale for clinical development of TMCM as a possible therapeutic to enhance resistance of the ocular mucosal barrier site against infectious disease, and points to its potential as an adjuvant for use in vaccination.

## STAR * METHODS

STAR★METHODS
| REAGENT or RESOURCE | SOURCE | IDENTIFIER |
| --- | --- | --- |
| **Antibodies** |  |  |
| FC: TCR$\gamma\delta$- APC (Clone: GL3) | eBioscience | 17-5711-82 (RRID: AB_842756) |
| FC: TCR$\alpha\beta$-FITC (Clone: H57-597) | BioLegend | 109205 (RRID: AB_314013) |
| FC: CD90.2- BV605 (Clone: 53-2.1) | BD | 536008 (RRID: AB_2665477) |
| FC: Vγ1- FITC (Clone: 2.11) | BioLegend | 141104 (RRID: AB_10697031) |
| FC: Vγ4- PE-CY7 (Clone: UC3-10A6) | eBioscience | 25-5828-82 (RRID: AB_2573474) |
| FC: Ly6C-BV510 (Clone: HK1.4) | BioLegend | 128033 (RRID: AB_2563363) |
| FC: Ly6G- APC-CY7 (Clone:1A8) | BioLegend | 127624 (RRID: AB_10640819) |
| FC: CD11b-BV785 (Clone: M1/70) | BioLegend | 101243 (RRID: AB_2562473) |
| FC: CD45-FITC (Clone: 30-F11) | BioLegend | 304006(RRID: AB_314275) |
| FC: IL-17A- BV650 (Clone: 17B7) | BD | 564170 (RRID: AB_2738641) |
| FC: CD44-BV786 (Clone: IM7) | BD | 563736 (RRID: AB_2738395) |
| FC: CD27-BV605 (Clone:LG.3A10) | BD | 563365 (RRID: AB_2738160) |
| FC: Phospho-p44/42 MAPK (Erk1/2) (Thr202/Tyr204) (197G2) | Cell Signaling | 98168S (RRID: AB_331773) |
| FC: Mouse anti-phospho-p38 MAPK (Clone: 28B10) | Cell Signaling | 9216 (RRID: AB_331296) |
| FC: RORγt-PE (Q31-38) | Biolegend | 129908(RRID: AB_2563456) |
| FC: CD69- BV785(Clone:H1.2F3) | BioLegend | 104543 (RRID: AB_2629640) |
| FC: Ki67- eFluor 450 (Clone: SolA15) | eBioscience | 69-5698-82 (RRID: AB_2637482) |
| FC: ViaKrome 808 Fixable Viability Dye | Beckman Coulter | C36628 (RRID: AB_2562423) |
| FC : CD3-BV510 (Clone: 17A2) | Biolegend | 100245 (RRID: AB_2562423) |
| FC: Myc-Tag (9B11) Mouse mAb | Cell Signaling | 2276 (RRID: AB_331726) |
| FC: IL-7R-PE (Clone: SB/199) | Biolegend | 121112 (RRID: AB_493509) |
| FC: CD121(IL1R)-APC | BioLegend | 113509 (RRID: AB_2264757) |
| FC: CD80-PerCP-Cy 5.5 (Clone: 16-10A1) | BD Biosciences | 560528 (RRID: AB_1727514) |
| FC: CD86-PE (Clone: PO3.1) | Biolegend | 12-0861-83 (RRID: AB_313159) |
| *In vitro*: Anti-mMincle | Invivogen | mabg-mmcl-2 (RRID: AB_2891969) |
| *In vitro*: Anti-IL-1R1 | BioXCell | BE0256 (RRID: AB_2661843) |
| *In vivo*: Anti-TCR$\gamma\delta$ (Clone: UC7) | BioXCell | BE0070 (RRID: AB_1107751) |
| *In vitro*: Anti-IL-6R | BioXCell | BE0047 (RRID: AB_1107588) |
| *In vitro*: Anti-TNF-a | BioXCell | BE0070 (RRID: AB_1107751) |
| **Bacterial and virus strains** |  |  |
| *Pseudomonas aeruginosa* | Shanks Lab (Pitt) | https://pubmed.ncbi.nlm.nih.gov/11581056/ |
| *Corynebacterium mastitidis* | Caspi Lab | https://pubmed.ncbi.nlm.nih.gov/28709803/ |
| *Corynebacterium glutamicum* | ATCC | 13032 |
| *Pks/FadD32::Tn* | St. Leger Lab | This paper |

| Chemicals, peptides, and recombinant proteins |  |  |
| --- | --- | --- |
| CaCl<sub>2</sub> | Quality Biological | 351-130-721 |
| Collagenase D | Roche | 45-11088866001 |
| DNase I | Roche | 50-100-3290 |
| Dispase II | Gibco | 17-105-041 |
| Liberase<sup>TM</sup> | Sigma | 5401119001 |
| Brefeldin A (Golgi-Plug) | BD PharMingen | 555029 (RRID: AB_2869014) |
| PMA/Ionomycin (Lymphocyte Activation Cocktail) | BD PharMingen | 550583 (RRID: AB_2868893) |
| Perm/Wash Buffer | BD Biosciences | 554723 |
| BD Phosflow<sup>TM</sup> Fix Buffer I | BD Biosciences | 557870 |
| Recombinant human IL-1β | R&D system | 201-LB |
| Methanol Optima LCMS grade | Fisher Scientific | A4564 |
| Water Optima LCMS grade | Fisher Scientific | W64 |
| Acetonitrile Optima LCMS grade | Fisher Scientific | A9554 |
| Isopropanol Optima LCMS grade | Fisher Scientific | 4614 |
| Chloroform ACS grade | Fisher Scientific | C606SK4 |
| Acetic Acid Optima LCMS grade | Fisher Scientific | A11350 |
| Hexanes | Fisher Scientific | H302SK-4 |
| Ammonium acetate | Fisher Scientific | A11450 |
| Aminopropyl SPE Columns | Waters Corp | WAT020840 |
| C18 SPE Columns | Waters Corp | WAT020805 |
| Silica SPE Columns | Waters Corp | WAT054930 |
| Silica TLC plates (10x10 cm) | Millipore Sigma | 1.05631.0001 |
| Primuline | Millipore Sigma | 206865-1G |
| Butylated Hydroxytoluene | MP Biomedicals | 101162 |
| Acetone | Fisher Scientific | A949SK-4 |
| Corynomycolate (α18:1/my18:1) | Avanti Polar Lipids | Custom synthesis |
| Trehalose monocorynomycolate (α18:1/my18:1) | Avanti Polar Lipids | Custom synthesis |
| Trehalose dicorynomycolate (α18:1/my18:1, α18:1/my18:1) | Avanti Polar Lipids | Custom synthesis |
| Ethanol | Sigma | 24112 |
| Corn oil | Sigma | C8267 |
| Tamoxifen | Sigma | T5648 |
| LB Broth, Miller | Fisher Scientific | BP1426-2 |
| Agar powder/flakes | Fisher Scientific | BP1423-2 |
| Fosfomycin | Fisher Scientific | 50-136-8271 |
| Tween-80 | Sigma | P4780 |
| Crystal violet | Fisher Scientific | C581-25 |
| Corning Falcon test tube with cell strainer snap cap | Corning | 352235 |
| 5mm stainless steel beads | Qiagen | 69989 |
| EZ-Tn5 Transposase | Lucigen | TNP92110 |
| Isoflurane | Baxter | NDC: 10019-360-60 |
| Ketamine | VET ONE | NDC: 13985-584-10 |
| Xylazine | Dechra | NDC: 17033-099-05 |
| Tropicamide | Bausch & Lomb | NDC 24208-585-59 |
| Phenylephrine | Bausch & Lomb | NDC 82260-103-05 |

| Critical commercial assays |  |  |
| --- | --- | --- |
| CD11c MicroBeads UltraPure, mouse | Miltenyi | 130-125-835 |
| LEGENDplex™ Mouse Inflammation Panel (13-plex) | BioLegend | 740446 |
| ELISA MAX™ Deluxe Set Mouse IL-17A | BioLegend | 432504 |
| Deposited data |  |  |
| TCR sequence data | This paper | https://dataview.ncbi.nlm.nih.gov/object/PRJNA1226517?reviewer=im4d6m5dge01k98anlbmk2p2qq |
| Experimental models: Organisms/strains |  |  |
| C57BL/6J | Jackson Lab | JAX 000664. |
| TLR2-/- | Jackson Lab | JAX 004650 |
| Nur77<sup>GFP</sup> | Jackson Lab | JAX 016617 |
| Knock-in CD3ζ switch mice | Paul Love Lab<br>https://www.nichd.nih.gov/research/atNICHD/Investigators/love | https://pubmed.ncbi.nlm.nih.gov/37945821/ |
| Software and algorithms |  |  |
| FlowJo v10.4 | Tree Star | https://www.flowjo.com |
| GraphPad Prism 9.1.1 | GraphPad | https://www.graphpad.com |
| VDJFasta | Centivax Inc. | https://github.com/immunoengineer/vdjfasta |
| SciexOS 3.1.6.44 | AB Sciex | https://sciex.com/products/software/sciex-os-software |
| Other |  |  |

## CONTACT FOR REAGENT AND RESOURCE SHARING

Further information and requests for resources and reagents should be directed to and will be fulfilled by the Lead Contact, Anthony St. Leger & Benjamin Schwarz and Rachel Caspi

### Data and Code availability

TCR sequence data for Reviewers’ access: https://dataview.ncbi.nlm.nih.gov/object/PRJNA1226517?reviewer=im4d6m5dge01k98anlbmk2p2qq. Public link will be provided upon acceptance of the manuscript.

## EXPERIMENTAL MODEL AND SUBJECT DETAILS

### Mice

Inbred wild-type (WT) C57BL/6J, TLR2-/-, and Nur77^GFP^ mice on the C57BL/6 background were obtained from Jackson Labs. Knock-in CD3ζ switch mice^19^ were provided by Dr. Paul E. Love. All mice were housed in specific pathogen-free conditions at National Eye Institute (NEI) or the University of Pittsburgh. Care and use of animals followed institutionally approved animal study protocols and Animal Research Advisory Committee (ARAC) guidelines.

## METHOD DETAILS

### Tissue harvesting and processing

Mice were euthanized by CO2 followed by extensive cardiac perfusion with PBS. Two conjunctivae of one mouse were harvested by excising the eyelid and bulbar conjunctiva, pooled, minced, and digested in Liberase^TM^ (50 ug/ml) containing medium for 45 min at 37℃ with shaking. After Liberase^TM^ treatment, conjunctivae were separately filtered through a 70 μM filter by crushing and homogenizing using a 1 cc syringe plunger.

### *In vivo* tamoxifen treatment

Knock-in CD3ζ switch mice ^19^ were orally gavaged with 200 µl of a 10 mg/ml tamoxifen suspension in corn oil daily for 5 consecutive days. Switch efficiency from 6Y to 6F was assessed by intracellular staining with anti-Myc 3 days after the last gavage.

### Cell purification and culture

CD11c+ cells were purified from splenocytes using CD11c MicroBeads UltraPure (MACS, Miltenyi Biotech). γδ T cells were obtained by staining lymphocytes with fluorescently labeled anti-CD90.2 and anti-TCRγδ, followed by Fluorescence-Activated Cell Sorting (FACS).

CD11c+ cells (1×10^5^) were co-cultured with TCRγδ+ cells (1×10^4^) in 200 μl RPMI supplemented with 10% FBS for 48 to 72 hours under indicated conditions. Brefeldin A (Golgiplug) was added in the last 6 hrs of culture. Culture supernatants were assessed for cytokine abundance by ELISA (R&D Systems) and cells were stained for γδTCR, Ki67, IL-17 to assess intracellular IL-17 by flow cytometry. Where noted, Mincle, IL-1R, TNF-α, and IL-6R were inhibited with antibodies such as anti-mMincle IgG(10 µg/ml) anti-IL1R (1 µg/ml), anti-TNF-α (10 µg/ml), and anti-IL-6R (10 µg/ml), respectively.

### Crude lipid fractionation by Solid-Phase Extraction

For all organic and solid phase extractions (SPE) liquid chromatography mass spectrometry (LCMS) grade, or highest available purity, solvents and methanol washed glass vessels were used to ensure against contamination. Sham extraction samples were taken through all steps to ensure against the introduction of artifacts at any step.

Starting from a frozen bacterial pellet of approximately 1-gram wet weight 4 ml of methanol was added and the sample agitated to suspend the pellet. Subsequently, 5 ml of water and 5 ml of chloroform was added and the sample agitated to facilitate a Bligh & Dyer biphasic extraction. The sample was centrifuged at 10,000 g for 20 minutes to enforce layering. The bottom, organic phase, was retained and dried under nitrogen at 60°C. The organic fraction, referred to as the crude lipid, was resuspended in 2 ml of chloroform for further characterization.

The *C. mast* crude lipid and subsequent fractions were analyzed by thin layer chromatography on a silica plate eluted with a chloroform:methanol:water mixture in a 7:3:0.25 volumetric ratio. Lipid spots were resolved by spraying with Primuline (10 mg/ml in acetone) and imaged under UV light.

In an effort to separate the phospholipids from the corynomycolates a dual phase SPE strategy was employed ^36^. An aminopropyl cartridge was placed upstream of a silica cartridge and the system was conditioned with 5 ml of 1:1 hexanes:chloroform v/v and eluted via vacuum. An aliquot of the resuspended crude lipid (500 µL) in chloroform was loaded onto the columns and washed with an additional 10 ml of chloroform. The flow through and initial chloroform wash were collected as fraction 1(SPE1_F1). A 10 ml volume of 4.5:4.5:1 hexanes:isopropanol:chloroform v/v + 0.1 % glacial acetic acid was applied to the column and eluent collected as fraction 2 (SPE1_F2). A 10 ml volume of 3:6:1 v/v hexanes:isopropanol:chloroform:methanol v/v + 0.2 % glacial acetic acid was applied and collected as fraction 3 (SPE1_F3). A 10 ml volume of 0.5:8:0.5:1 v/v hexanes:isopropanol:chloroform:methanol was applied and collected as fraction 4 (SPE1_F4). Lastly a 10 ml volume of 2:8 v/v chloroform:methanol + 50 mM ammonium acetate was applied and collected as fraction 5 (SPE1_F5). All fractions were dried under nitrogen at 60°C. Fractions were subsequently resuspended in 6:1 isopropanol:methanol with 5 µg/ml butylated hydroxytoluene (BHT) and analyzed via TLC as described above before proceeding to mass spec analysis.

Following determination of activity and initial MS characterization, SPE1_F3 and SPE1_F4 were combined and subjected to orthogonal SPE separation aimed at fractionation of free fatty acids and mycolic acids. The combined fractions were diluted 1:1 in 1:3 v/v methanol:water to increase polarity of the solution for loading. A C18 SPE cartridge was conditioned with 20 ml 1:3 v/v methanol:water. Samples were loaded and chased with an additional 10 ml volume of 1:3 v/v methanol:water with the flow through collected as fraction 1 (SPE2_F1). A 10 ml volume of 75% acetonitrile in water was applied and collected as fraction 2 (SPE2_F2). A 10 ml volume of acetonitrile was applied and collected as fraction 3 (SPE2_F3). Lastly a 10 ml volume of chloroform was applied and collected as fraction 4 (SPE2_F4). All fractions were dried as above and either tested for activity or resuspended in 6:1 isopropanol:methanol with 5 ug/ml BHT for analysis by MS.

### Mass Spectrometry

All MS analysis was performed on a 7500 QTrap instrument (AB Sciex). All liquid chromatography was performed on a LD-40XR series HPLC. Initial analysis of crude lipid extract or SPE fraction was pursued via direct injection in both negative and positive polarity. High intensity peaks were selected visually and fragmentation spectra were collected using a ramping of collision energy. For targeted profiling of corynomycolates and free fatty acids, a series of multiple reaction monitoring (MRM) ion pairs were established using a conserved fragmentation logic for each family of molecules (Supp table). All targets assayed were detected using negative polarization. Only targets shown to be present in direct injection assays were included. Free fatty acids were detected using a neutral loss of 44 m/z fragment air for all species detected. Unconjugated corynomycolates were detected using a conserved product anion for the α-chain of the molecule i.e. -281.2 m/z for the a18:1 series. Both TMCM and TDCM species were detected via the m/z -323.1 trehalose-associated fragment. Synthetic corMyc(α18:1/my18:1), TMCM(α18:1/my18:1), and TDCM(α18:1/my18:1, α18:1/my18:1) (Avanti Polar Lipids) were utilized to calibrate the method. Lipids were eluted using a Synergi 4 um Fusion-RP 80Å (50 x 2 mm) column (Phenomenex) held at 50°C with gradient from 1:1 v/v acetonitrile:water 5 mM ammonium acetate to 5 % v/v water in acetonitrile 5 mM ammonium acetate over 5 minutes at a flow rate of 0.75 ml/min. Integration values for eluted peaks were normalized to the total sum of the free fatty acid signal to assess the enrichment of corynomycolate species in each fraction. All LCMS data processing was done using SciexOS 3.1.6.44 (AB Sciex).

### Generation of *C. mast* mutants

A *C. mast* Tn mutant library was created as described previously^20^. Briefly, an EZ-Tn5 Transposase (Lucigen, Middleton, WI, USA) was incorporated with genes for mCherry and Kanamycin resistance. This transposon construct was incorporated with *C. mast* electrocompetent cells, and electroporated using a Gene Pulser Xcell Total System (Bio-Rad, Hercules, CA, USA) at 1.8 kV, 25 µF, and 200 ohms. Bacteria were then recovered in liquid LBT and placed in a 37°C shaking incubator for 1-2 hours. After recovery, bacteria were plated on Kanamycin selective LBT plates. The corMyc deficient *Pks/FadD32::Tn* mutant was isolated and identified as described ^20^

### Characterization of mutant *C. mast*

#### Biofilm formation

A high throughput screen for biofilm formation was done by using a crystal violet based assay^37^. Bacteria were grown in a 96 or 48 well tissue culture treated dish in a 37°C incubator for 48 hours. After incubation, growth was quantified at OD600, supernatants were removed, and wells were washed three times with DI water. Biofilm was then stained with 100 μl of 0.1% crystal violet and incubated at room temperature for 15 minutes. After incubation, wells were washed three times with DI water, and 100 μl of 190 proof ethanol was added to the well and incubated at room temperature for 15 minutes. Plate was then analyzed by quantifying absorbance in a plate reader at 550 nm. The amount of biofilm was normalized to growth by dividing amount of biofilm by amount of growth.

### Application of bacteria to the ocular surface

Four-week-old WT C57BL/6J mice lacking *C. mast* were first swabbed with a sterile cotton swab to disrupt the tear film. Eyes were then inoculated with 5x10^5^ CFU in 5 μl of PBS every other day for a total of three inoculations. Ocular colonization was assessed one week after final inoculation by swabbing the conjunctiva with a sterile swab, and plating on LBT plates with Fosfomycin.

### Assessment of *in vivo* immunity

Three weeks after final inoculation, mice were sacrificed by cardiac perfusion and cervical draining lymph nodes (DLNs) and conjunctivae were isolated. Conjunctivae were isolated as previously described^4,20^. Briefly, eyelids and bulbar conjunctivae were excised, pooled from both eyes, minced, and incubated in a collagenase solution at 37°C for one hour. DLNs were also minced and incubated in a collagenase solution at 37°C for 30 minutes. After incubating, conjunctivae and DLNs were filtered through a 40-µm filter using the plunger from a 3-cc syringe plunger. Samples were then washed two times with RPMI + 10% FBS. Conjunctival samples were then filtered using a Corning Falcon Test Tube with Cell Strainer Snap Cap (352235; Corning Inc., Corning, NY, USA). Two thirds of the conjunctival samples were then stained for immune cell neutrophil analysis, or together with the DLN samples were stimulated with phorbol myristate acetate (PMA)/ionomycin in the presence of brefeldin A for 4 hours at 37°C. After the 4 h incubation, samples were stained for Vγ4 TCR and IL-17A. All samples were analyzed using CytoFLEX LX Flow Cytometer and FlowJo v10.8.

### Optical coherence tomography measurement

Mice were anesthetized using Ketamine/Xylazine combination, and images of cornea were acquired by anterior segment Optical Coherence Tomography (OCT) with 10-mm telecentric lenses by Bioptigen Spectral Domain Ophthalmic Imaging System Envisu R2200 (Bioptigen Inc., Durham, NC).

### P. aeruginosa challenge

Four-week-old WT C57BL/6J mice lacking *C. mast* were inoculated with *C. mast* or *Pks/FadD32::Tn* as described above, or every day for two weeks with synthetic lipids. 24 hours after the final lipid inoculation, mice were challenged with *P. aeruginosa* as described previously^4,34^. The next day, whole eyes were collected, homogenized in sterile PBS using 5 mm stainless steel beads (Qiagen, Hilden, Germany) and the MP FASTPREP 24 (MPBio, Santa Ana, CA). Homogenized eyes were then serially diluted and plated on LB agar. After overnight culture at 37°C, number of colonies were quantified.

### Reconstitution of synthetic lipids

Previously frozen 5 mg aliquots of synthetic corynomycolate(α18:1/my18:1), trehalose monocorynomycolate(α18:1/my18:1), trehalose dicorynomycolate(α18:1/my18:1, α18:1/my18:1) powders in glass vials were brought to room temperature, and dissolved in either 1 ml of ethanol (for *in vitro* experiments) or 1ml of corn oil (for *in vivo* experiments) and incubated at 37°C water bath with shaking for 1 hour until fully dissolved. Dissolved lipids were stored at 4°C for 1-week storage and at -20°C for long-term storage.

### Stimulation of reconstituted lipids *in vitro* and *in vivo*

Bring glass bottles containing dissolved lipids to room temperature. For *in vitro* experiments, 5 µg/ml of each lipid (dissolved in ethanol) were added to the DC-γδ T co-culture and incubated for 2 days at 37°C.

For *in vivo* experiments, mice were first anesthetized in isoflurane. Then 2 µl of indicated lipid (5 mg/ml) dissolved in corn oil were instilled into the upper fornix pocket and 2 µl into the bottom fornix pocket of each mouse. Mice were kept undisturbed in the isoflurane chamber for 5-10 minutes before returning to their cages.

### ELISA

Culture supernatant was collected from DC cultures or DC- γδ T cocultures with indicated stimulation. Biolegend ELISA MAX™ Deluxe Set Mouse IL-17A (Cat# 432504), or LEGENDplex™ Mouse Inflammation Panel (13-plex) were used according to manufacturer’s protocols.

### Single-cell γδ TCR sequencing

Single-cell suspension was prepared from cervical LNs of C57BL/6J mice with either corn oil or crude lipids and were stained with antibodies against αβTCR, γδTCR, Vγ4, CD44, and CD27. Single cells with αβTCR- γδTCR+Vγ4+CD44highCD27low phenotype were sorted into 96-well PCR plates. RNA isolation, cDNA library preparation, and sequencing were performed as described^38^. The resulting sequences were analyzed by VDJFasta as described previously^15,39^.

### Statistical analysis

Data are presented as mean ± SEM. Mann-Whitney test, or Brown-Forsythe and Welch ANOVA, or one-way ANOVA, or Welch’s T test were used to calculate p-values as indicated in Figure legends using GraphPad Prism v10.

## ACKNOWLEDGMENTS

This work was supported by the intramural Research Program of the NIH, National Eye Institute (project EY000184; R.R. Caspi), R01EY032482 (A. J. St. Leger), U24EY035102 (A. J. St. Leger), P30EY008098, T32EY017271 (Y. E. Rigas), Research to Prevent Blindness (New York, NY) (A. J. St. Leger), the Eye and Ear Foundation of Pittsburgh (A. J. St. Leger), and the Division of Intramural Research/NIAID/NIH (Research and Technologies Branch; B. Schwarz). Preliminary method development, leading to the LCMS methods and data displayed here, was supported by the Immunity to Pulmonary Pathogens Section and Catharine M. Bosio within the NIIAD intramural program.

The authors sincerely thank the National Eye Institute and National Heart Lung and Blood Institute Flow Cytometry Core facilities for assistance in cell sorting and the Visual Function Core (NEI) for assistance in OCT measurements. We thank Dr. Yueh Hsiu Chien (Department of Microbiology and Immunology, Stanford University School of Medicine, Stanford) for her guidance in single-cell TCR sequencing.

Graphical abstract was created with BioRender.com.

## AUTHOR CONTRIBUTIONS

XX and YR planned, performed and analyzed experiments and interpreted data. XX wrote the first manuscript draft. JG performed scRNA-Seq and wrote the original code; JG, VN, AZ and TJ analyzed and interpreted scRNA-Seq data. GG and PEL provided the CD3 switch knock-in mice and discussed data. MC, AG, BX, ZP, EB, CR, NTB, JS and YJ performed experiments; MTM and MC evaluated and interpreted *in vivo* ocular surface data; MJM interpreted data, provided counseling on experimental design, reviewed and edited the manuscript. RRC, BS and AJSL conceptualized the study design and approaches and edited the manuscript. RRC conceived and supervised the study, reviewed and finalized the manuscript. All co-authors reviewed and approved the manuscript.

## Supplementary Figures

**Figure S1.**
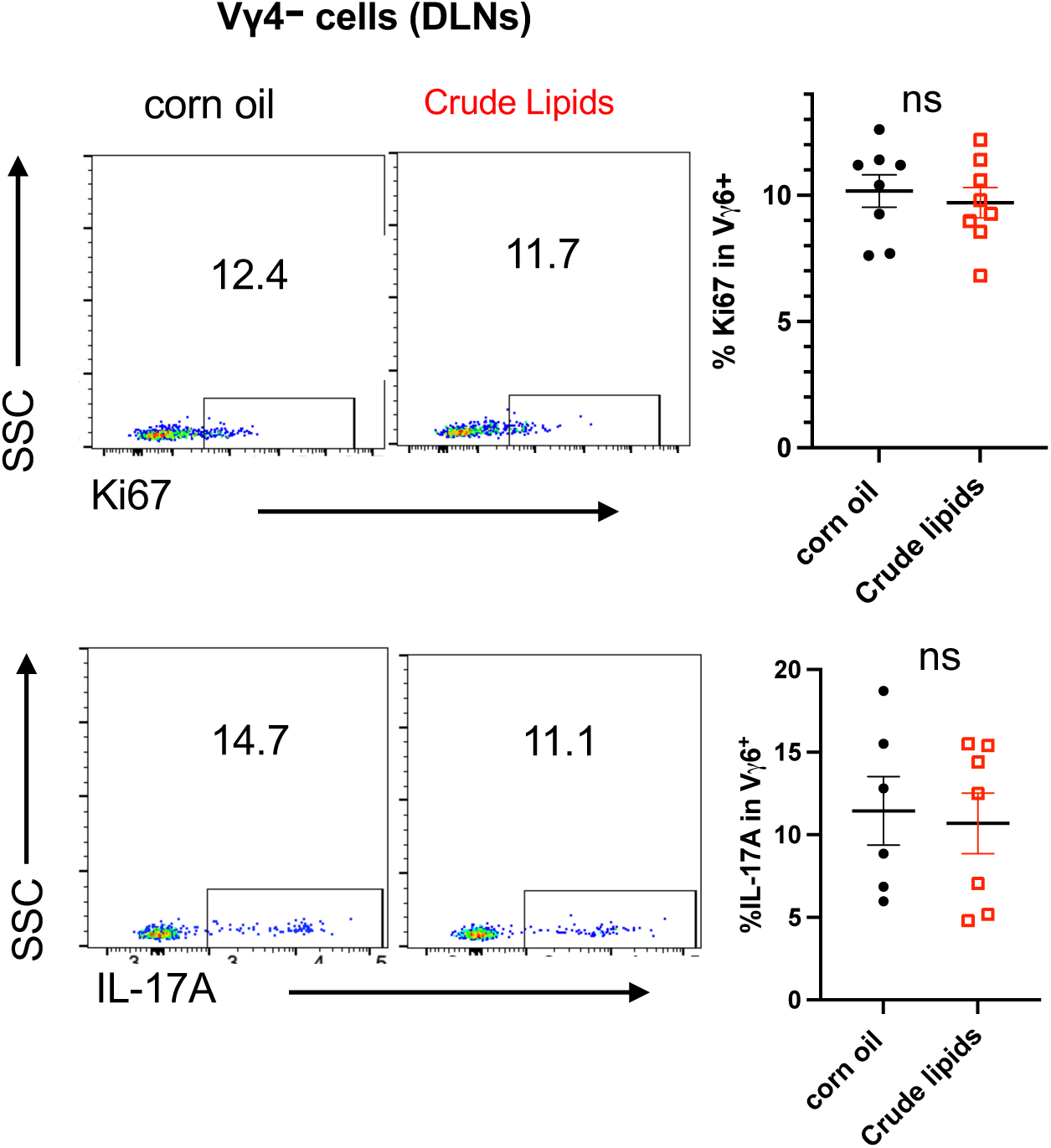
Lipid components of C. mast do not elicit ΙL-17 responses in Vγ4-negative cells (related to Figure 1) Representative flow cytometry plots and dot plots showing the percentage of Ki67+ and IL-17A+ in Vγ4- cells of DLNs from mice described in Figure 1C. Each dot represents one animal. Bars represent mean ± SEM. *p<0.05, **p<0.01, ***p<0.001. Statistical significance was determined by the Mann-Whitney test.

**Figure S2.**
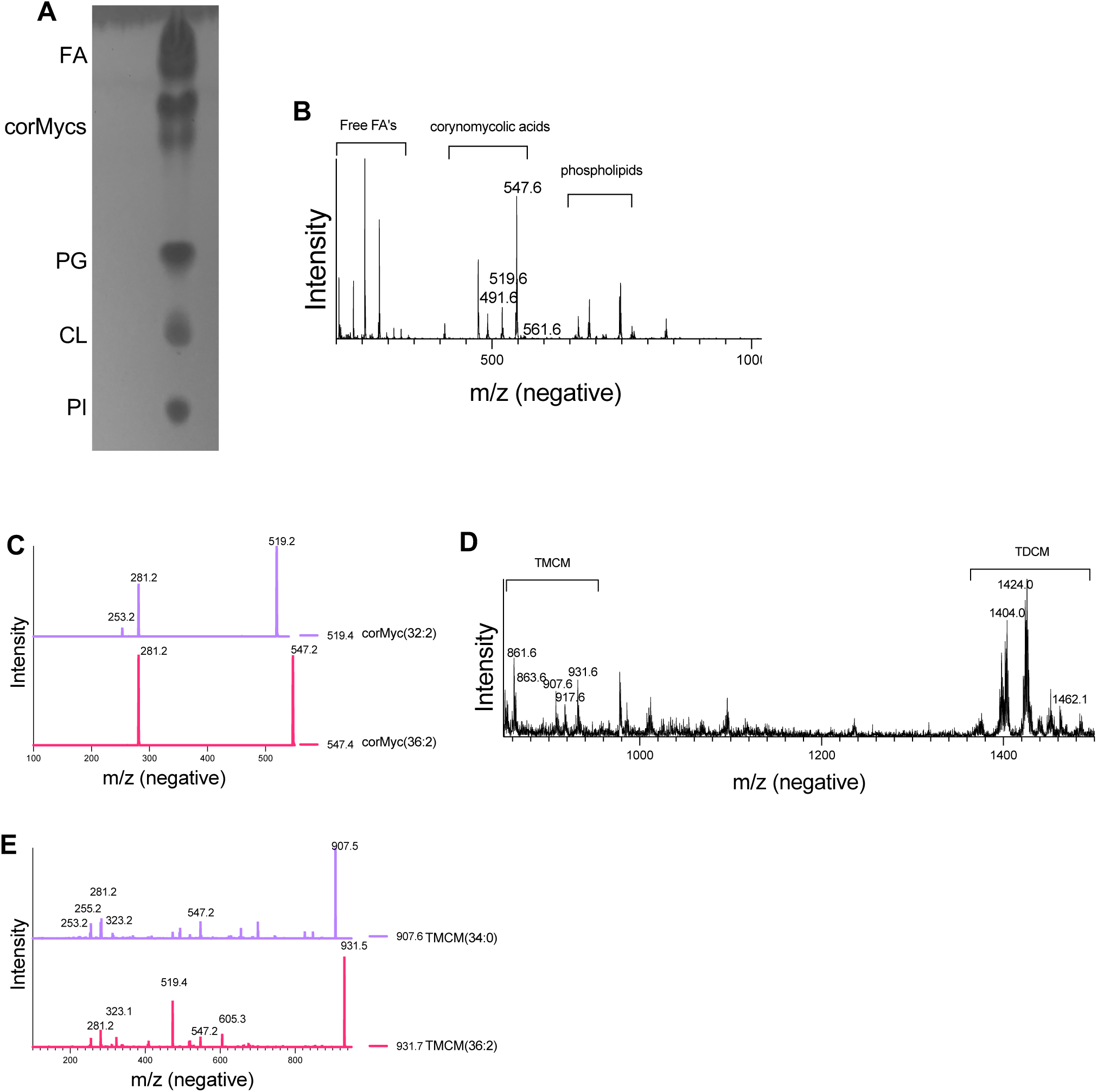
Identification and characterization of lipid classes in *C. mast* crude lipid extract (related to Figure 2) (**A**) Crude lipids of *C. mast* were analyzed by thin layer chromatography(TLC). (**B**) Full mass spectra under negative ionization conditions of crude lipid preparation from WT *C. mast*. 200-1100 m/z regions were populated by free fatty acids, corynomycolic acids(corMycs), and phospholipids(PG,CL,PI). Within the corynomycolic acid region, major peaks are anticipated to be corMyc(C32:2) at -519.6 (M-), corMyc(C36:2) at -547.6 (M-). (**C**) Fragmentation spectra under negative polarity of the putative corMyc (32:2) at -519.6 and corMyc(C36:2) at -547.6 using rolling collision energy establishing these species as corMyc(a18:1/my18:1) under the -547.4 signal, indicated by the -281.2 product ion, and a mixture of corMyc (a16:1/my16:1) and corMyc (a18:1/my14:1) contributing to the -519.2 ion, indicated by the -253.2 and –281.2 product ions respectively. (**D**) Zoomed spectra from the same injection as **(B)** for 850 -1500 m/z region of trehalose conjugated corynomycolates in WT *C.mast* crude lipid preparation. Putative acetate series (M+59-) for TMCM and TDCM species are labeled. The major isomeric contributors to each signal are anticipated to be TMCM (C34:0) at -907.5 (M+AcO-), and TMCM(C36:2) at -931.5 (M+AcO-), TDCM(72:4) at -1462.1 (M+AcO-), TDCM (68:4) at -1404.0 (M+AcO-). (**E**) Fragmentation spectra of putative acetate series ions (M+59-) for TMCM(C34:0) at -907.5 (M+AcO-), and TMCM(C36:2) at -931.5 (M+AcO-) showing a distribution of species under each signal as evidenced by the series of fatty acid peaks in the 200-300 m/z fragment range.

**Figure S3.**
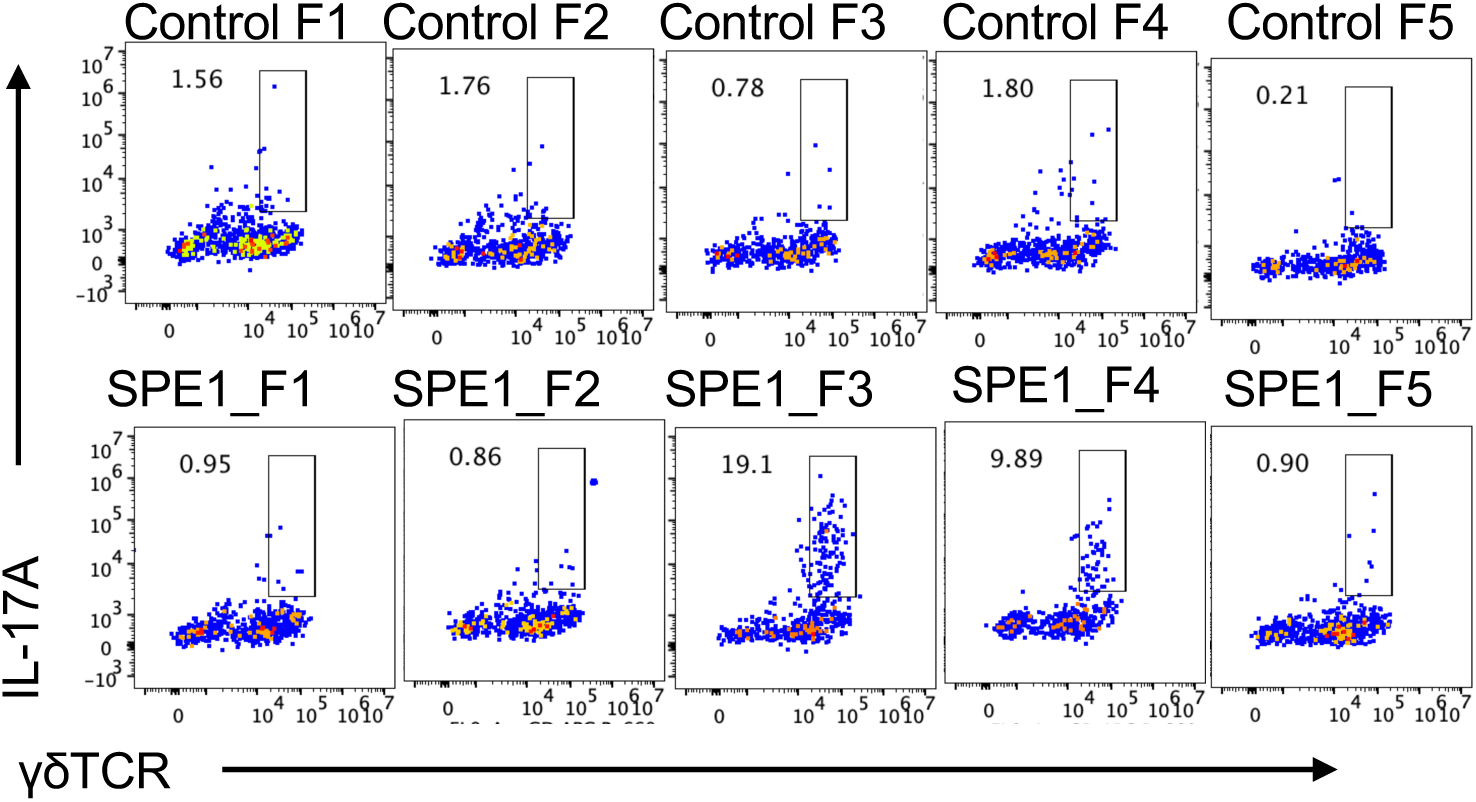
SPE1 fractions F3 and F4 contain il-17 stimulatory activity (related to Figure 2) Representative FACS plots showing the percentage of IL-17A in the co-cultures described in Figure 2D.

**Figure S4.**
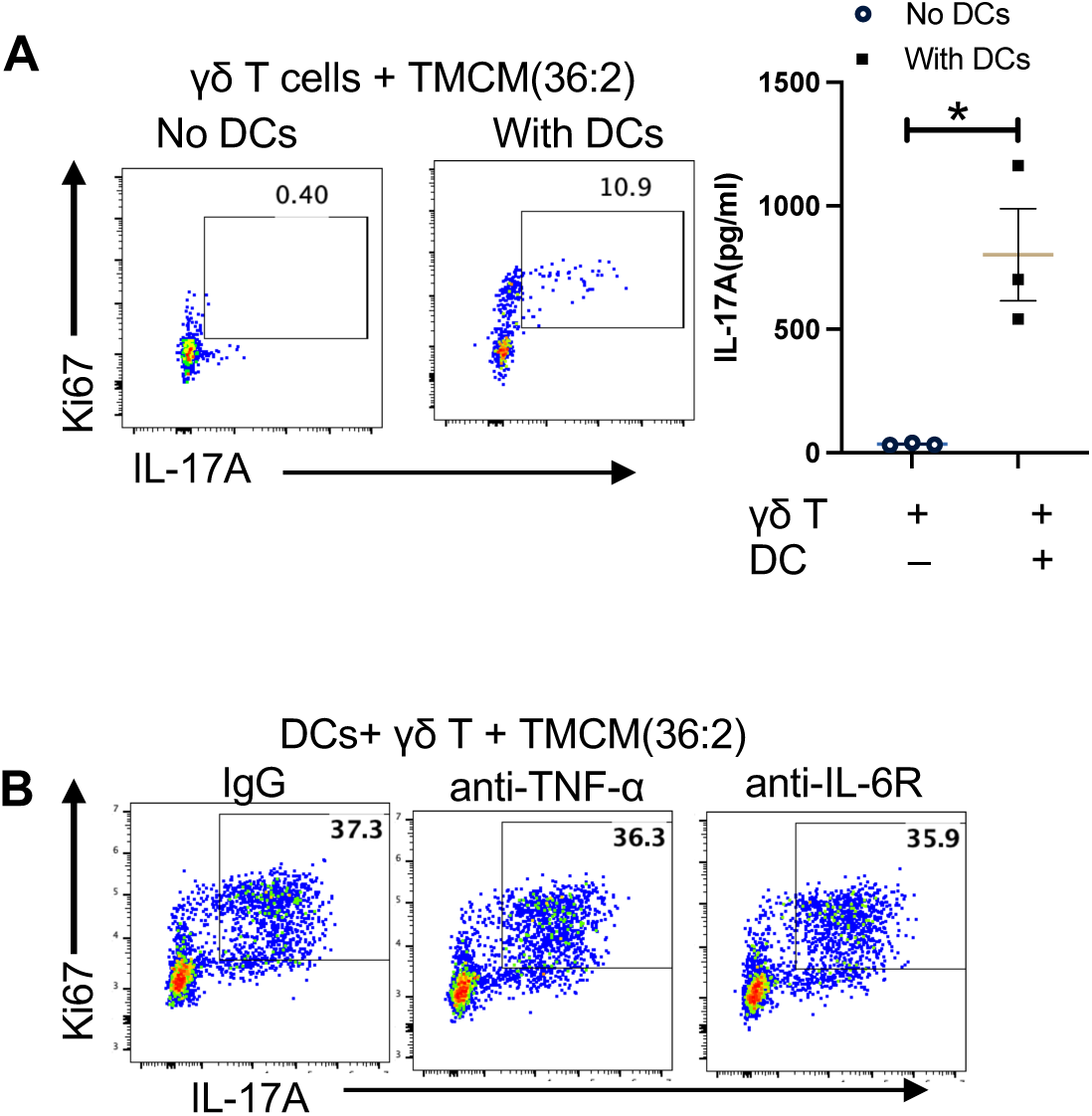
TNF-α and IL-6 affect TMCM-induced IL-17 responses when present together, but not individually (related to Figure 4) (**A**) Sorted γδT cells alone or with CD11c+ DCs in the presence of TMCM(36:2) for 3 days. Representative flow cytometry plot showing the percentage of Ki67+IL-17A+ on gated γδT cells, and levels of IL-17A in the culture supernatant are shown in dot plot. Each dot represents one experiment. (**B**) Blockade antibodies targeting anti-TNF-a or anti-IL-6R were added in cocultures with sorted γδT cells with CD11c+ DCs in the presence of TMCM(36:2) for 2 days. Representative flow cytometry plot showing the percentage of Ki67+IL-17A+ on gated γδT cells. Representative data of at least 2 independent experiments. Bars represent mean ± SEM. *p<0.05, **p<0.01, ***p<0.001. Statistical significance was determined by the Mann-Whitney test (A).

**Figure S5.**
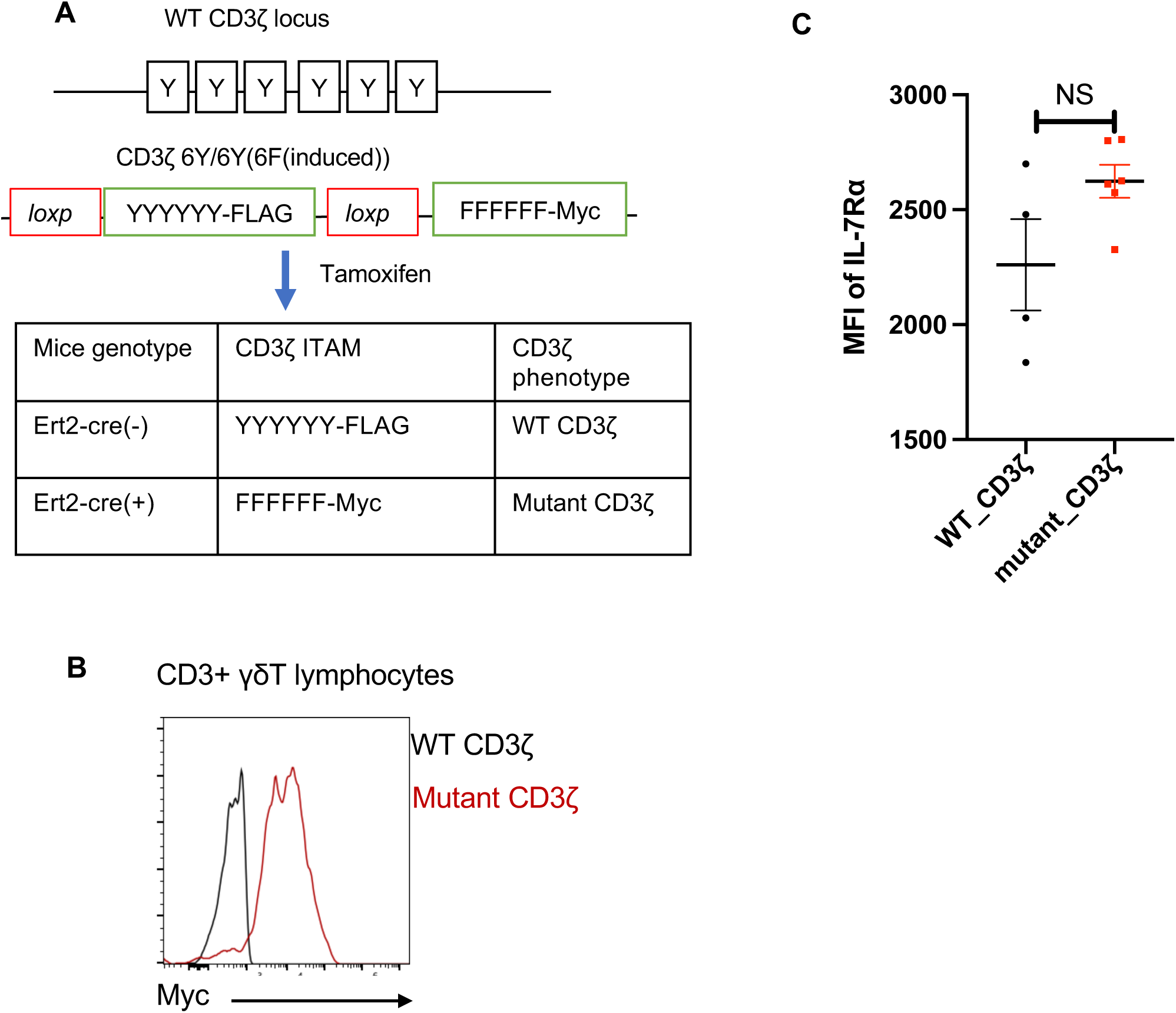
γδ TCR signaling does not affect IL-7R expression (related to Figure 5) (**A**) Schematic representation of CD3ζ ITAM motif in Ert2-cre(-) and Ert2-cre(+) CD3ζ 6Y/6Y(6F(induced)) mice before and after tamoxifen induced modification. The tyrosines (Y) in the wild-type ITAM motif were replaced by phenylalanines(F) in mutant CD3ζ, impairing TCR signaling. (**B**)The efficiency of mutant CD3 induction shown as Myc tag expression after tamoxifen treatment. Lymphocytes from WT and mutated CD3ζ mice stained with anti-Myc. (**C**) The expression of IL-7Rα on CD44+CD27- γδ T cells from WT and mutated CD3ζ mice 8 days after tamoxifen treatment. Each dot represents an individual mouse. The results were representative of at least 2 independent experiments. Bars represent mean ± SEM. *p<0.05, **p<0.01, ***p<0.001.Statistical significance was determined by Mann-Whitney test (C).

**Figure S6.**
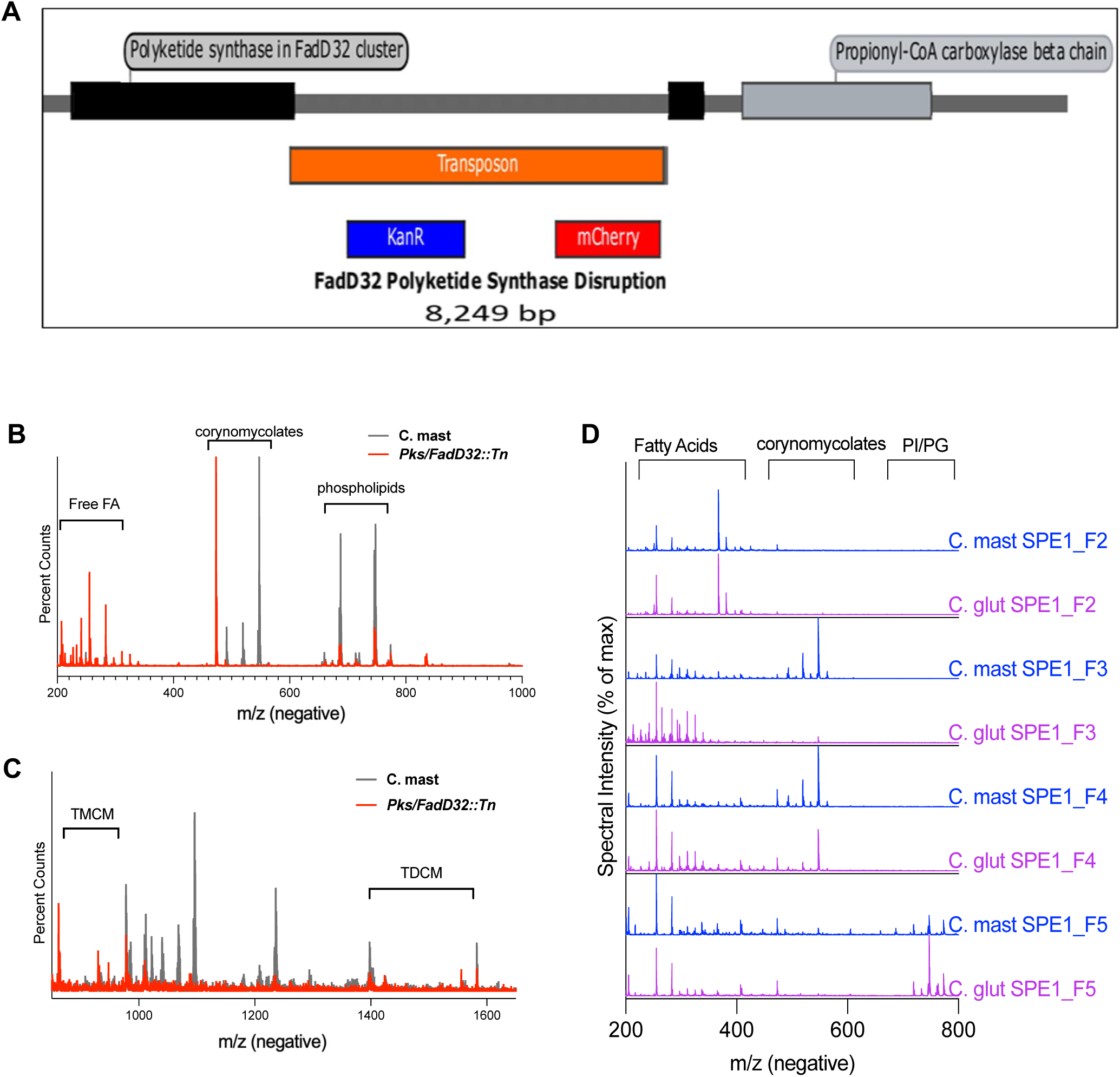
*C. mast Pks/FadD32::Tn* lacks corynomycolates expressed by WT *C. mast* and *C. glut (*related to Figure 6) **(A)** The whole genome of *Pks/FadD32::Tn* was sequenced, and the transposon (Tn) was inserted in FadD32 cluster polyketide synthase. **(B-C)** Full mass spectra under negative ionization conditions of crude lipid preparation from WT *C. mast Pks/FadD32::Tn* mutant *C. mast*. 200-1000 m/z regions were populated by free fatty acids, corynomycolates (corMycs), and phospholipids (**B**). Zoomed spectra for 850 -1500 m/z region displayed trehalose conjugated corynomycolates in WT and *Pks/FadD32::Tn* mutant *C. mast* crude lipid preparation **(C)**. **(D)** Lipid fractions from *C. mast* and *C. glut* lipidome were examined by liquid chromatography-mass spectrometry (LC-MS). F1 showed a redundant pattern to F2 and is not depicted.

**Figure S7.**
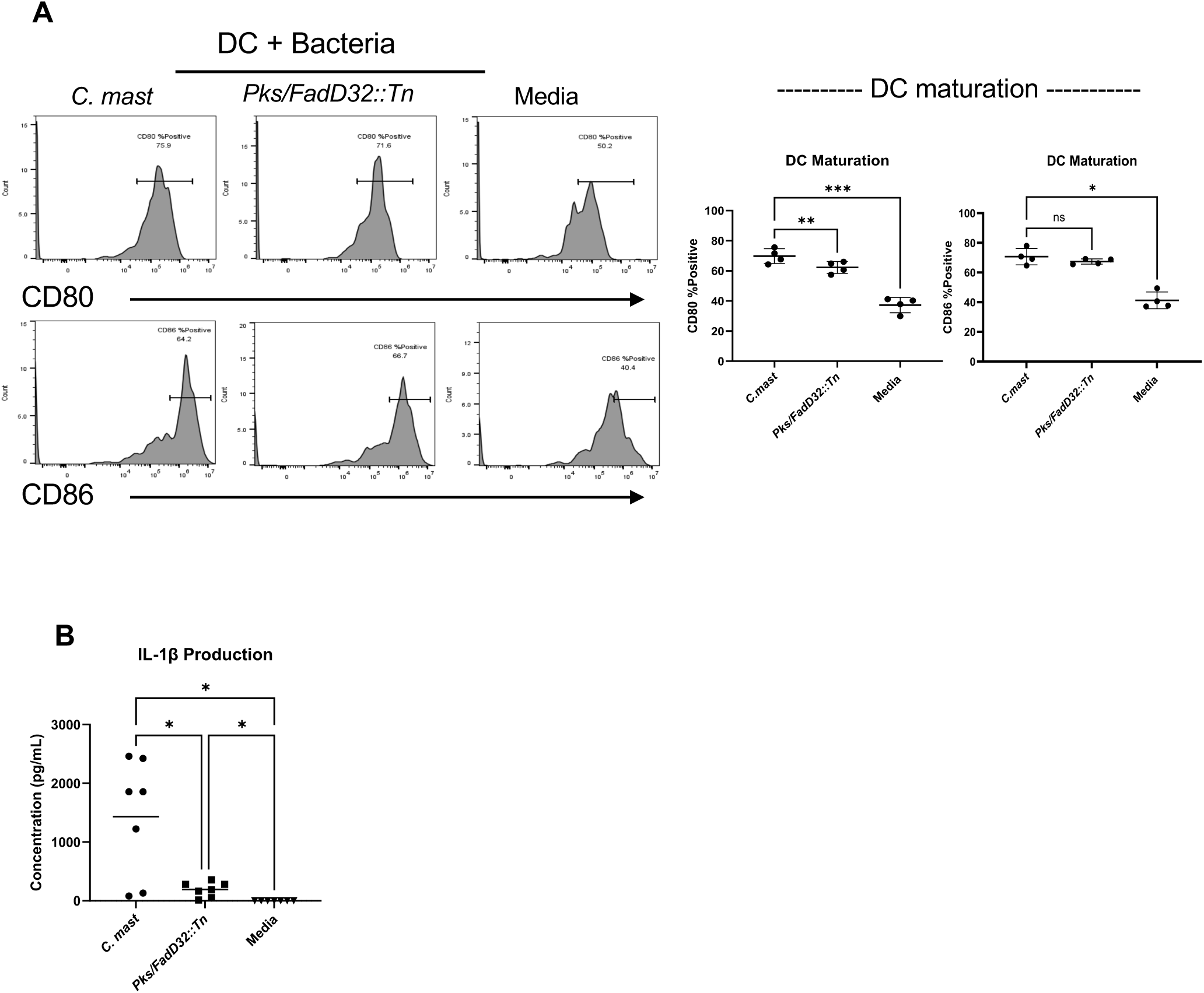
*Pks/FadD32::Tn* mutant *of C. mast* induces a reduced DC IL-1β response (related to Figure 6) **(A-B)** 5x10^4^ BMDCs were co-cocultured with 1x10^5^ CFU of either WT *C. mast* or *Pks/FadD32::Tn*. After incubating for 48 hours, BMDCs were assessed for DC maturation through expression of CD80 and CD86 (**A)**, and supernatants were analyzed for IL-1β by ELISA (**B**). In all graphs, bars represent mean ± SEM. Symbols represent individual experiments. Significance was determined using paired Brown-Forsythe and Welch ANOVA (A) or one-way ANOVA (B) (*p= 0.0049, ***p= 0.0001).

**Figure S8.**
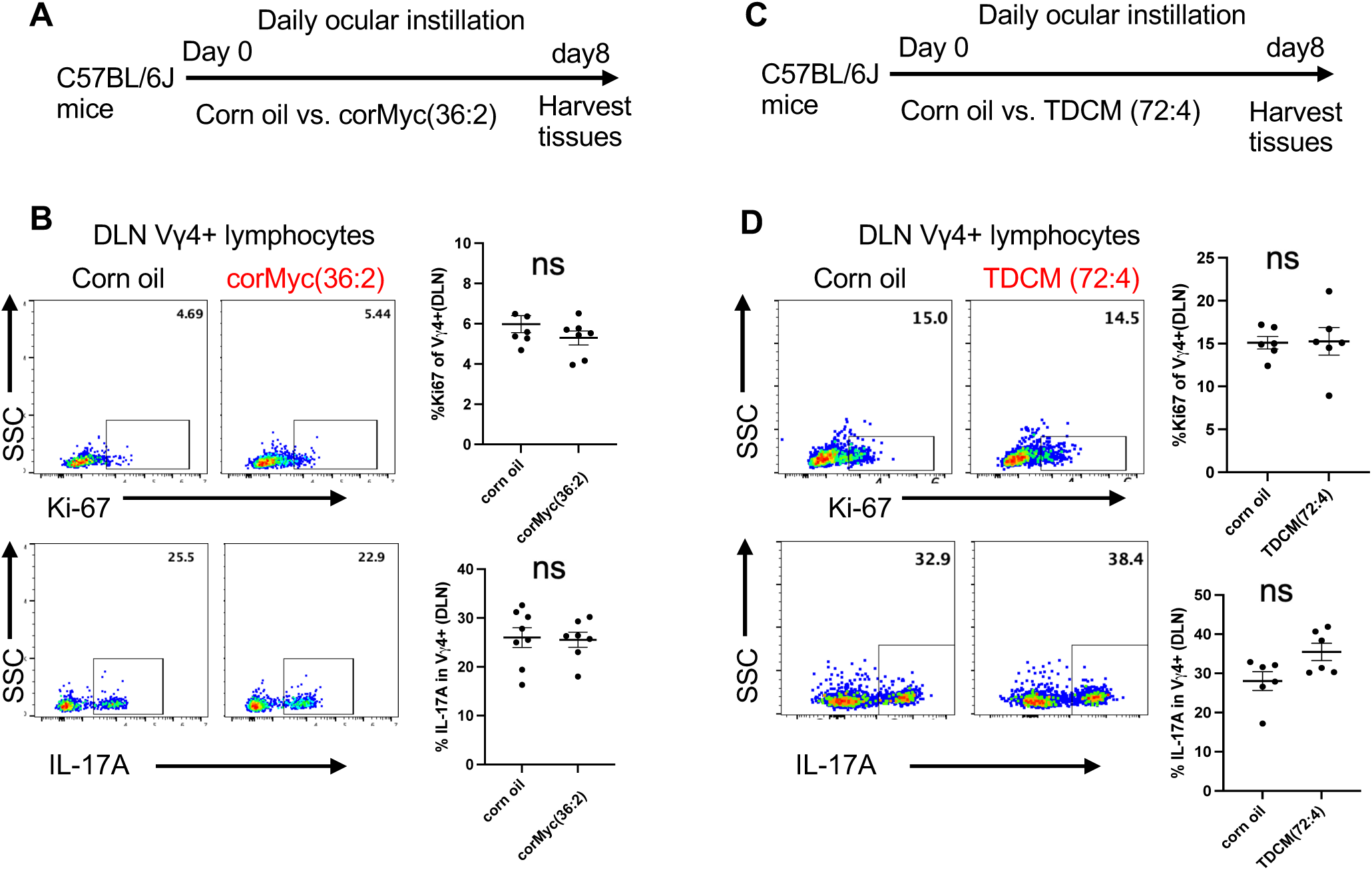
corMyc (36:2) and TDCM (72:4) do not elicit ocular IL-17 responses in healthy mice (related to Figure 7) (**A-B**) Experimental design to assess ocular immune responses elicited by synthetic corMyc (36:2). (**A**) Corn oil (vehicle control) or 25 µg synthetic corMyc (36:2) was instilled on the ocular surface of C57BL/6J mice daily for 7 days. (**B**) Representative flow cytometric analysis and dot plots show the percentage of Ki67+ or IL-17A+ Vγ4 cells in the eye-draining cervical LNs (DLNs). Each dot represents one animal. (**C-D**) Experimental design to assess ocular immune responses elicited by synthetic TDCM(72:4). (**C**) Corn oil (vehicle control) or 25 µg synthetic TDCM(72:4) was instilled on the ocular surface of C57BL/6J mice daily for 7 days. (**D**) Representative flow cytometric analysis and dot plots show the percentage of Ki67+ or IL-17A+ Vγ4 cells in the eye-draining cervical LNs (DLNs). Each dot represents one animal. The results were representative of at least 2 independent experiments. Bars represent mean ± SEM with *p<0.05, **p<0.01, ***p<0.001. Statistical significance was determined by Mann-Whitney test. ns, not significant (**B,D**).

